# Behavioural variability and cortical electrophysiological signals depend on recent outcomes during human reinforcement motor learning

**DOI:** 10.1101/2021.04.29.441455

**Authors:** Patrick Wiegel, Meaghan Elizabeth Spedden, Christina Ramsenthaler, Mikkel Malling Beck, Jesper Lundbye-Jensen

**Affiliations:** Department of Nutrition, Exercise and Sports, University of Copenhagen, Copenhagen, Denmark; Department of Sport Science, University of Freiburg, Freiburg, Germany; Klinik für Palliativmedizin, Universitätsklinikum Freiburg, Freiburg, Germany; Wolfson Palliative care Research Centre, Hull & York Medical School, University of Hull, Hull, UK; Cicely Saunders Institute, Department of Palliative care, Policy & Rehabilitation, King’s College London, London, UK; Department of Neuroscience, University of Copenhagen, Copenhagen, Denmark

**Keywords:** Reinforcement, motor learning, prefrontal cortex, variability, neural oscillations, exploration and exploitation.

## Abstract

The history of our actions and the outcomes of these represent important information, which can inform choices, and efficiently guide future behaviour. While unsuccessful (S-) outcomes are expected to lead to more explorative motor states and increased behavioural variability, successful (S+) outcomes lead to reinforcement of the previous action and thus exploitation. Here, we show that during reinforcement motor learning, humans attribute different values to previous actions when they experience S- vs. S+ outcomes. Behavioural variability after S- outcomes is influenced more by the previous outcomes compared to what is observed after S+ outcomes. Using electroencephalography, we show that neural oscillations of the prefrontal cortex encode the level of reinforcement (high beta frequencies) and reflect the detection of reward prediction errors (theta frequencies). The results suggest that S+ experiences ‘overwrite’ previous motor states to a greater extent than S- experiences and that modulations in neural oscillations in the prefrontal cortex play a potential role in encoding the (changes in) movement variability state during reinforcement motor learning.

## Introduction

A striking feature of the nervous system is its ability to plan motor actions, to monitor the consequences during and following motor actions and to integrate these in future actions. During repeated practice of a given task, unsuccessful (S-) outcomes of our actions require exploration of new task solutions, while successful (S+) actions should ideally be reproduced and the strategy should be exploited. This process is called reinforcement learning (RL) and it relies on the evaluation of performance outcomes (Sutton & Barto, 1998).

RL guides behavioural variability in subsequent actions and optimises future performance. The level of reinforcement can be estimated by measuring changes in trial-to-trial variability (TTV) of behavioural characteristics (e.g. kinematic parameters). Although TTV has long been considered and in some ways can be an unwanted by-product of a noisy sensorimotor system, it is also an important aspect of motor learning (Wu *et al*., 2014). TTV may be separated into unintended variability (due to noise) and intended variability (due to exploration). Provided that the outcome is monitored and evaluated by the central nervous system, variability may indeed serve to guide learning processes towards those behavioural characteristics, that lead to desirable outcomes. In this way, changes in TTV across time are necessary components of motor learning. During motor learning, TTV is increased after failures but decreased after rewards and this points towards a reward-dependent modulation of TTV to maximise future rewards (Takikawa *et al*., 2002; Galea *et al*., 2013; Pekny *et al*., 2015). However, the outcome of the current action is not the only information used to guide subsequent behaviour. Instead, findings in rodent experiments demonstrate that the past several outcomes can be integrated to control future movements in rodents (Dhawale *et al*., 2019), but in humans, knowledge on the regulation of TTV in response to previous outcomes during reinforcement motor learning is largely limited.

To which degree past outcomes are relevant during reinforcement motor learning may depend on the outcomes of – and thus the value attributed to - previously performed actions. Past actions are only informative when they help the agent to perform better in future trials. Previous S- actions help to delineate actions that should be avoided. As a consequence, other solutions can be tested in future actions, and S- outcomes can thus guide exploration, potentially leading to S+ outcomes. When an S+ outcome is experienced, this should lead to exploitation. But are past motor actions also helpful or merely disregarded when experiencing S+ outcomes during reinforcement motor learning? In this case, information about earlier movements might be less valuable or down-weighted since the agent naturally aim to reproduce the current S+ movement. Nevertheless, history of previous actions may still inform future actions. Here, we tested this assumption and designed a reinforcement motor learning task in which human participants performed goal-directed wrist flexion movements. We measured TTV in wrist angle at movement end point and investigated influences of different previous outcomes as a measure of the level of reinforcement.

Previous studies hypothesised that reward-dependent learning is mediated by the difference between expected and actual rewards, so-called reward prediction errors (Schultz, 2017). These signals drive learning based on feedback on outcome and serve as the basis for future behavioural adjustments. A variety of neural circuits in subcortical and cortical systems have been implicated in this context (Watanabe, 1996; Knutson *et al*., 2000; Schultz, 2000; Schall *et al*., 2002; Cohen *et al*., 2007; Histed *et al*., 2009; Narayanan *et al*., 2013; HajiHosseini & Holroyd, 2015; Levy *et al*., 2020). Lately, a growing body of evidence suggests that neural circuits of the prefrontal cortex contribute to reward-based learning in monkeys (Watanabe, 1996; Kim & Shadlen, 1999; Barraclough *et al*., 2004; Seo & Lee, 2008; Histed *et al*., 2009) and humans (Akitsuki *et al*., 2003; Cohen *et al*., 2007; Marco-Pallares *et al*., 2008; HajiHosseini *et al*., 2012; HajiHosseini & Holroyd, 2015). Modulations of neural oscillations over frontal cortical areas can be observed after outcome information in decision-making tasks potentially reflecting outcome-guided learning (Luft, 2014). Specifically, increases in oscillatory activity have been observed for S- outcomes in theta band frequencies (4 – 8 Hz) and for S+ outcomes in high beta/low gamma frequencies (25 – 35 Hz). Neural oscillations have been suggested to subserve important functions for the regulation of information transfer across the brain and for controlling synaptic plasticity during learning (Buzsaki & Draguhn, 2004; Luft, 2014). Thus, modulations of oscillatory activity during outcome-processing constitute a reasonable mechanism to drive adjustments in behaviour during reinforcement motor learning. Although the role of neural oscillations during feedback-based learning is well described in cognitive (decision-making) tasks, modulations of neural oscillations during human reinforcement motor learning are not well understood. To test whether different behavioural outcome scenarios are associated with specific oscillatory reinforcement signals in the prefrontal cortex during motor learning, we recorded electroencephalography (EEG) while participants practiced the motor task.

Thus, in the present paper we tested the effects of different outcomes and history of previous actions on TTV and oscillatory signals in the prefrontal cortex during human reinforcement motor learning. We hypothesised that S+ actions would result in behavioural reinforcement i.e. exploitation and reduced consideration of previous outcomes. In contrast, we assumed that S- actions would lead to greater behavioural exploration that is informed by and thus depends more on previous outcomes. In addition, we hypothesised that S+ outcomes would lead to increased oscillatory activity in high beta frequencies and S- outcomes to increased oscillatory activity in theta frequencies in the prefrontal cortex.

## Results

Twenty-six participants performed wrist flexion reaching movements to horizontally move a computer cursor into a target area (Fig. 1a & 1b). Movements were guided by a visual scene on the computer screen which informed participants about movement outcomes (Fig. 1c). During the experiment, participants were confronted with different target positions and they received no online feedback but only binary augmented feedback on the outcome (i.e. S+ or S-) following each trial. Movement end points for a representative participant during the main protocol are plotted in Fig. 1d. Participants showed greater movement end point variability during the main protocol than during performance of movements to a stationary target with visual online feedback (Fam1, F-test, F = 319.5, P < 0.001) and without visual online feedback (Fam2, F-test, F = 239.7, P < 0.001, Fig. 1e; Fam2 vs. Fam1: F-test, F = 22.2, P < 0.001).

**Figure 1.**
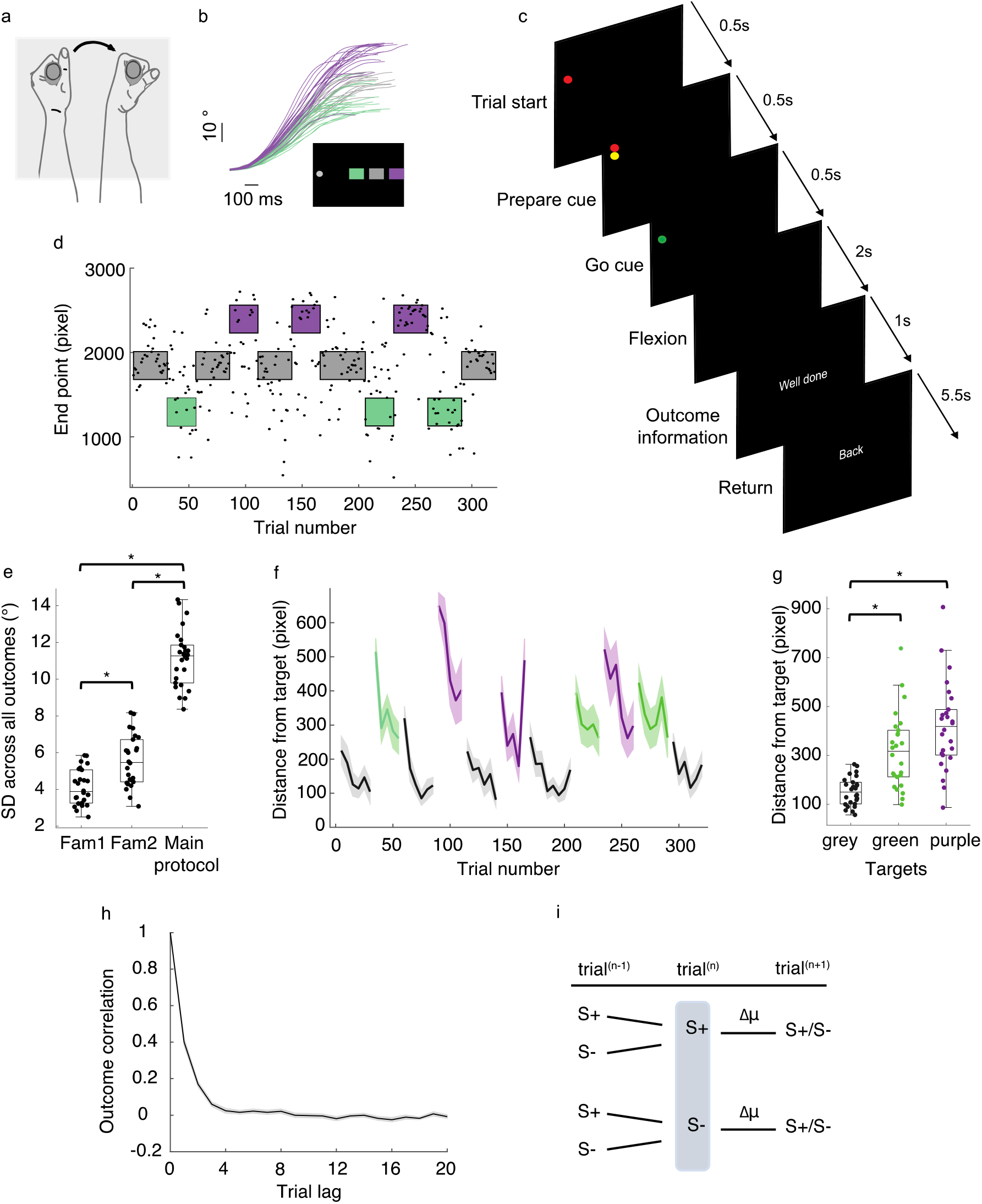
Motor variability during reinforcement motor learning. **a** Participants grabbed a handle to perform discrete wrist flexion movements with their left hand. The sight of the hand and forearm was hidden by a custom-made box (grey shaded area) to remove visual feedback of the moving arm. **b** The wrist angle was recorded by a goniometer that was integrated into the handle. The panel shows 60 exemplary wrist angle traces from a participant aiming at the green, grey and purple target box (20 movements each). Data were aligned to movement start and movement end. The rectangle illustrates the computer cursor and the approximate positions of the different targets. Cursor movements were one-dimensional in the horizontal plane. **c** The motor task was guided by different visual scenes on the computer screen. Trial start was indicated by a red light appearing on the upper left side of the computer screen. After 0.5 s, a yellow light below the red light signalled that participants had to prepare their movements. Finally, after another 0.5 s a green light indicated the Go cue. Participants had a time window of 2 s to perform their movements. Both, the cursor (representing the wrist angle) and the target area were invisible on the screen. Thus, after each trial participants received binary feedback about the outcome of the movement. S+ trials were indicated by “Well done” while S- trials were indicated by “Try again”. In both cases, feedback was visible for 1 s. After this period, participants were asked to move their wrist back to the starting position. A new trial started after 5.5 s. **d** Movement end points (in pixel) from one exemplary participant. Note that we changed the horizontal position of the target several times during the experiment (after 25, 30 or 40 trials) to stimulate exploring motor behaviour of the participants. **e** Boxplots and individual data for standard deviations from Fam1, Fam2 and the main protocol. * indicate significant differences between the conditions. Note that x-axis values were jittered to more clearly present the data. **f** The grand average data for distance from target (pixel) during the main protocol. Note that the figure shows averaged data in bins of 5 trials. Shading represents the standard error of the mean. **g** Boxplots and individual data for the distance from the different targets (in pixel) averaged across the main protocol. * indicate significant differences between the conditions. **h** We performed partial autocorrelation of the outcome time series (S+ and S- outcomes). The plot shows the grand averaged data. Shading represents the standard error of the mean. **i** Our framework focused on the question how the past two trials (trial^(n)^ and trial^(n-1)^) influence regulations of TTV (Δμ) in movement endpoint and oscillatory reinforcement signals.

In general, cursor end point distance from the targets varied considerably. The grand average data are illustrated in Fig. 1f. The data suggest that while participants approximated the targets positions quite well during blocks when the target did not move, the distance from the target became much greater when the target position changed. Also, participants were generally better at approximating the target they were familiarized with (grey target in Fam1 and Fam2) compared to the unknown targets (grey vs. green target: t = −5.8, P < 0.001; grey vs. purple target: t = −6.9, P < 0.001; green vs. purple target: t = −1.8, P = 0.080) (Fig. 1g).

To investigate the strength of the relationship between previous and future outcomes, we performed PAC for the outcome time series of all movements of the main protocol (Fig. 1h). The analysis showed that the association of previous outcomes (trial lags 1 – 20) with the current outcome (trial lag = 0) decays with increasing trial lags which is in line with a previous study in rodents (Dhawale *et al*., 2019). It also confirms our assumption that the past two outcomes have the greatest association with future outcomes. In the remainder of the paper, we focused on the impact of the previous two outcomes (trial^(n)^ & trial^(n-1)^) on TTV in the main protocol (Fig. 1i).

### TTV depends on the outcome of the previous movement

In a first step, we calculated the number of S+^(n)^ and S-^(n)^ movements for each participant. Table 1 summarizes the descriptive data of this analysis. There was no significant difference in the number of S+ and S- movements (paired t-test, t = −0.7, P = 0.499) suggesting that participants performed a similar number of S+ and S- trials. We also tested the effect of the previous outcome^(n)^ on motor performance in the following trial^(n+1)^. Participants had a greater proportion of S+^(n+1)^ trials (i.e. hits) when the preceding trial was S+^(n)^ (71,3%) compared to when the preceding trial was S-^(n)^ (28,1%) (Test of equal or given proportions, X²(1) = 1515.4, P < 0.001). This indicates better motor performance after S+^(n)^ trials compared to S-^(n)^ trials.

**Table 1.** *Grand average descriptive data (n=26) of the number of S+*^(n)^ *and S-*^(n)^ *events and the number of subsequent S+ events in trial^(n+1)^. SD = standard deviation, Min = minimum, Max = maximum*.

| | Grand average<br>number | Mean ± SD | Min | Max | Grand average number of S+ in<br>trial<sup>(n+1)</sup> |
| --- | --- | --- | --- | --- | --- |
| S+<sup>(n)</sup> | 3951 | 151.9 ± 30.0 | 94 | 216 | 2818 (71.3%) |
| S-<sup>(n)</sup> | 4161 | 160.0 ± 30.0 | 96 | 218 | 1168 (28.1%) |

Next, we analysed TTV in movement endpoint after S+ (Δμ| S+^(n)^) and S- (Δμ| S-^(n)^) trials. Fig. 2a and Fig. 2b show the grand average of the conditioned probability distributions. Visual inspection of the data suggested greater kurtosis after S-^(n)^ trials than after S+^(n)^ trials.

**Figure 2.**
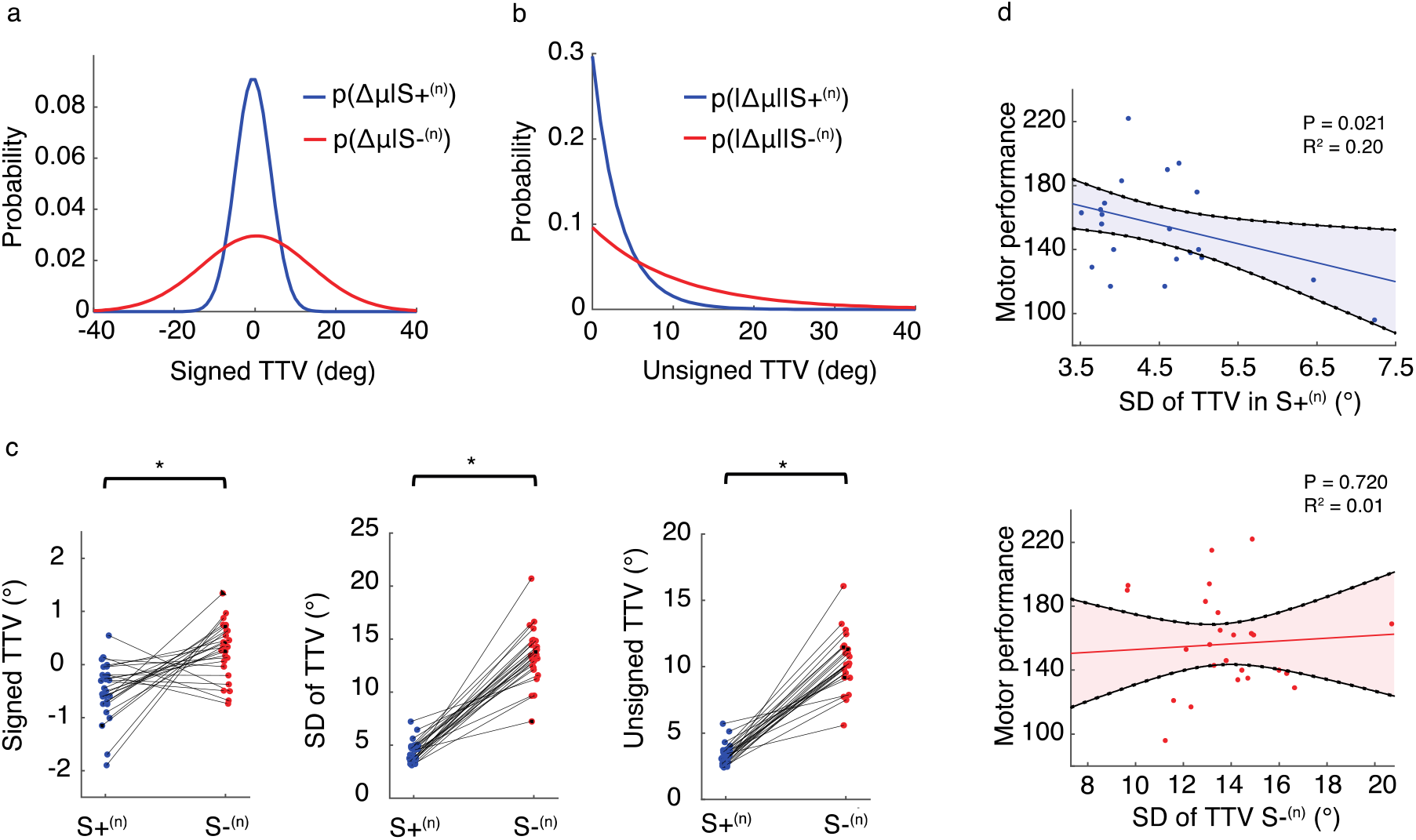
Behavioural variability depends on the previous outcome. **a** The grand average normal distribution for signed changes in TTV after S+^(n)^ and S-^(n)^ trials. **b** The grand average exponential distribution for unsigned changes in TTV after S+^(n)^ and S-^(n)^ trials. **c** Individual differences in signed TTV (left panel), SD of TTV (middle panel) and unsigned TTV (right panel) after S+^(n)^ and S-^(n)^ trials. * indicate significant differences between the conditions (after correction for multiple comparisons). Note that x-axis values were jittered to more clearly present the data. **d** Scatter plot showing the association between motor performance and SD of TTV after S+^(n)^ trials (top panel), and SD of TTV after S-^(n)^ trials (bottom panel). Each dot represents an individual participant. The lines represent the fitted regression line. Shading is 95% confidence intervals of the regression.

We tested this observation statistically and calculated the M (signed and unsigned TTV) and SD (signed TTV). The analyses supported our assumption from the visual inspection of the data and revealed significant differences in signed TTV (paired t-test, t = −3.6, P = 0.001, Fig. 2c left panel), the SD of TTV (F-test, F = 232.3, P < 0.001, Fig. 2c middle panel) and the unsigned TTV (F-test, F = 223.5, P < 0.001, Fig. 2c right panel). After S+^(n)^ movements, TTV in movement endpoint (signed TTV: −0.5° ± 0.6°; SD of TTV: 4.4° ± 1.2°; unsigned TTV: 3.4° ± 0.8°) was lower than after S- movements (signed TTV: 0.3° ± 0.5°; SD of TTV: 13.5° ± 2.6°; unsigned TTV: 10.4° ± 2.1°). The results indicate that variability of the movements and the absolute motor exploration were greater after S-^(n)^ movements than after S+^(n)^ movements. The differences in signed TTV suggest that participants tended to move the cursor slightly less after S+^(n)^ movements than after S-^(n)^ movements. Interestingly, differences in TTV were not only constrained to movement end point, rather TTV in maximal movement speed and movement time showed similar characteristics suggesting that participants reinforcement processes generalize to outcome-irrelevant parameters (Supplementary Fig. 1).

Moreover, SD of TTV in movement endpoint after S+^(n)^ movements was a good indicator of overall motor performance predicting the total individual score (i.e. total number of hits) (linear regression: F = 6.1, P = 0.021, R^2^ = 0.20, Fig. 2d top panel). This was not the case for SD of TTV in movement endpoint after S-^(n)^ movements (linear regression: F = 0.1, P = 0.720, R^2^ = 0.01, Fig. 2d bottom panel). Additional analyses demonstrated that the unsigned TTV after S+^(n)^ movements also predicted motor performance but not the signed TTV (Supplementary Fig. 2).

These results suggest different mechanisms of trial-by-trial reinforcement motor learning after S+^(n)^ compared to S-^(n)^ trials. S-^(n)^ trials stimulate greater TTV while S+^(n)^ trials lead to lower TTV. Moreover, participants who were able to “reproduce” S+ movements more accurately also performed better overall.

### The influence of previous outcomes differs for S+ and S- motor actions

It remains an open question whether the outcome of the second-to-last movement^(n-1)^ changes the impact of the outcome of the previous movement^(n)^. In other words, do outcomes prior to the current one have a different relevance when participants experience S+ ^(n)^ and S- ^(n)^ movements? To investigate this, we extracted TTV in movement endpoint conditioned on the previous two trials. Thus, we analysed TTV for four outcome scenarios: 1) S+^(n)^ & S+^(n-1)^, 2) S+^(n)^ & S-^(n-1)^, 3) S-^(n)^ & S+^(n-1)^ and 4) S-^(n)^_& S-_^(n-1)^.

The four conditions contained different trial numbers suggesting that certain outcome combinations were more likely than others. The rmANOVA with the number of trials as the dependent variable yielded a significant effect of outcome history (F_[1.3, 31.4]_ = 56.2, P < 0.001, η2_partial_ = 0.69). S+^(n)^ movements were more often preceded by S+^(n-1)^ movements than by S-^(n-1)^ movements (paired t-test: t = −9.2, P < 0.001). In contrast, S-^(n)^ movements were more often preceded by S-^(n-1)^ movements than by S+^(n-1)^ movements (paired t-test: t = 10.6, P < 0.001). The descriptive data of this analysis are shown in table 2.

**Table 2.** *Grand average descriptive data (n=26) of the number of* S+^(n)^ & S+^(n-1)^, S+^(n)^ & S-^(n-1)^, S-^(n)^ & S+^(n-1)^ and S-^(n)^ & S-^(n-1)^ events *and the number of subsequent S+ events in trial^(n+1)^. SD = standard deviation, Min = minimum, Max = maximum*.

| | Grand average<br>number | Mean ± SD | Min | Max | Grand average number of<br>$S^{+}$ in $\text{trial}^{(n+1)}$ |
| --- | --- | --- | --- | --- | --- |
| $S^{+(n)}$ & $S^{+(n-1)}$ | 2661 | $108.1 \pm 31.4$ | 48 | 177 | 1971 (75.1%) |
| $S^{+(n)}$ & $S^{-(n-1)}$ | 1136 | $45.5 \pm 7.8$ | 27 | 61 | 709 (62.4%) |
| $S^{-(n)}$ & $S^{+(n-1)}$ | 1145 | $45.2 \pm 7.6$ | 27 | 60 | 533 (46.6%) |
| $S^{-(n)}$ & $S^{-(n-1)}$ | 2910 | $116.2 \pm 31.8$ | 54 | 176 | 600 (20.1%) |

Next, we tested the effect of the different outcome histories on the proportion of S+ movements in trial^(n+1)^. A test of equal or given proportions revealed significant differences between the four outcome histories (X²(1) = 1691.9, P < 0.001) suggesting that motor performance depends on the past two outcomes. Indeed, the proportion of S+ movements in trial^(n+1)^ was highest in S+^(n)^ & S+^(n-1)^ (75.1%) and lowest in S-^(n)^ & S-^(n-1)^ (20.1%).

In a final step, we computed the TTV in movement endpoint for the different outcome histories. Due to the differences in trial number between the outcome histories, we performed bootstrapping with replacement (1,000 iterations) and matched the trial numbers for each participant. For each participant and condition, normal and exponential distributions were fitted from the bootstrapped datasets. The normal and exponential distributions for TTV after S+ trials^(n)^ conditioned on the trial^(n-1)^ are shown in Fig. 3a and Fig. 3b, respectively. These probability distributions appear to be very similar. As can be seen in Fig. 3c and Fig. 3d, this was different when trial^(n)^ was S-. The probability distribution after S- trials^(n)^ is broader when the preceding trial is also S-^(n-1)^ while it is narrower when the preceding trial is S+^(n-1)^. Statistical analyses were performed on the M and SD of the individual distributions. The results from the rmANOVA are presented in table 3.

**Figure 3.**
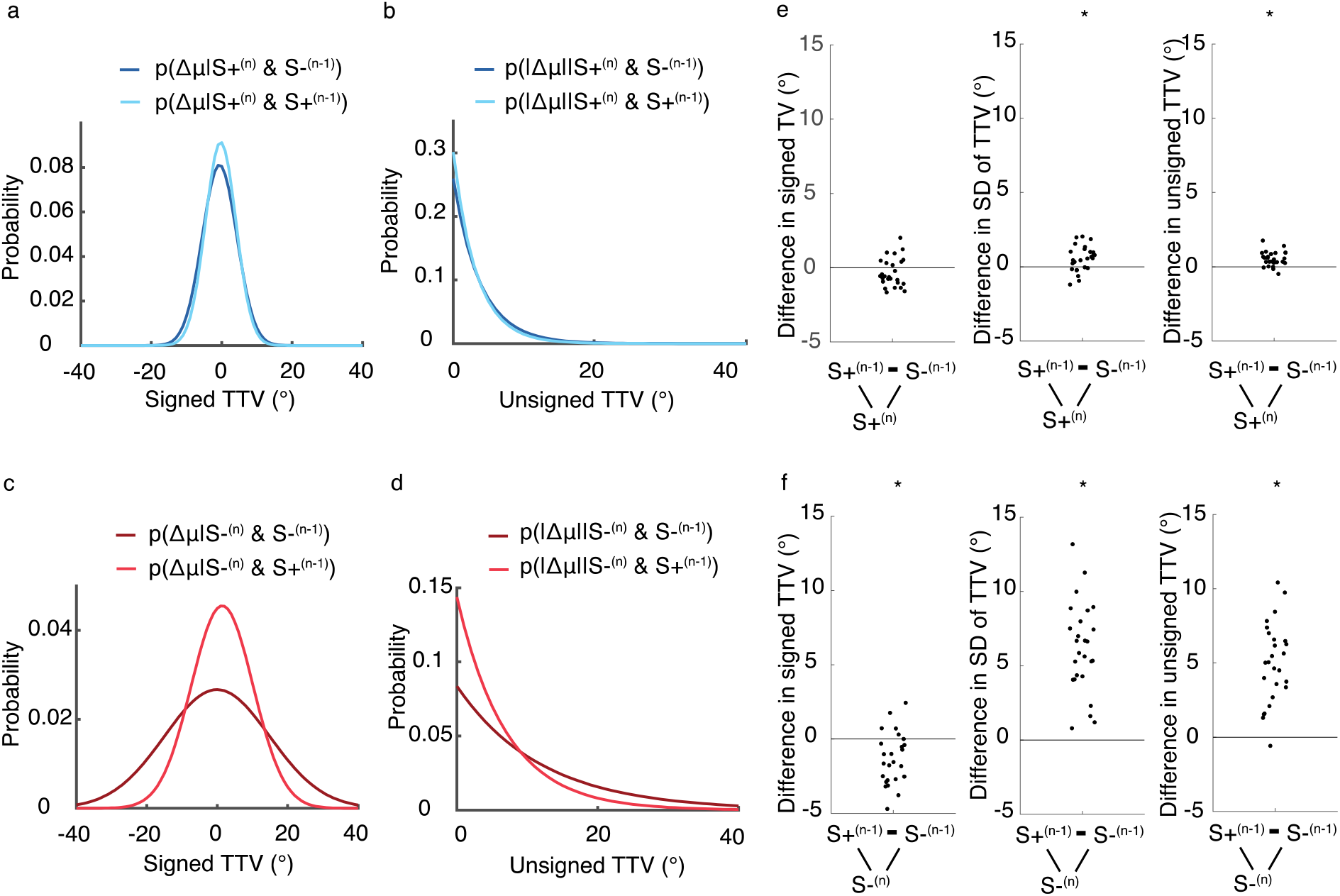
The impact of previous outcomes differs for S+ and S- motor actions. **a** The grand average normal distribution for signed TTV after S+^(n)^ trials when the previous trial was S+^(n-1)^ and S-^(n-1)^. **b** The grand average exponential distribution for unsigned changes in TTV after S+^(n)^ trials when the previous trial was S+^(n-1)^ and S-^(n-1)^. **c** The grand average normal distribution for signed TTV after S-^(n)^ trials when the previous trial was S+^(n-1)^ and S-^(n-1)^. **d** The grand average exponential distribution for unsigned changes in TTV after S-^(n)^ trials when the previous trial was S+^(n-1)^ and S-^(n-1)^. **e** Individual differences in signed TTV (left panel), SD of TTV (middle panel) and unsigned TTV (right panel) between S+^(n)^ that were preceded by S+^(n-1)^ and S-^(n-1)^ movements. **f** Individual differences in signed TTV (left panel), SD of TTV (middle panel) and unsigned TTV (right panel) between S-^(n)^ that were preceded by S+^(n-1)^ and S-^(n-1)^ movements. Each dot represents an individual. Note that x-axis values were jittered to more clearly present the data. * indicate significant differences between the different outcome histories (after correction for multiple comparisons).

**Table 3.**
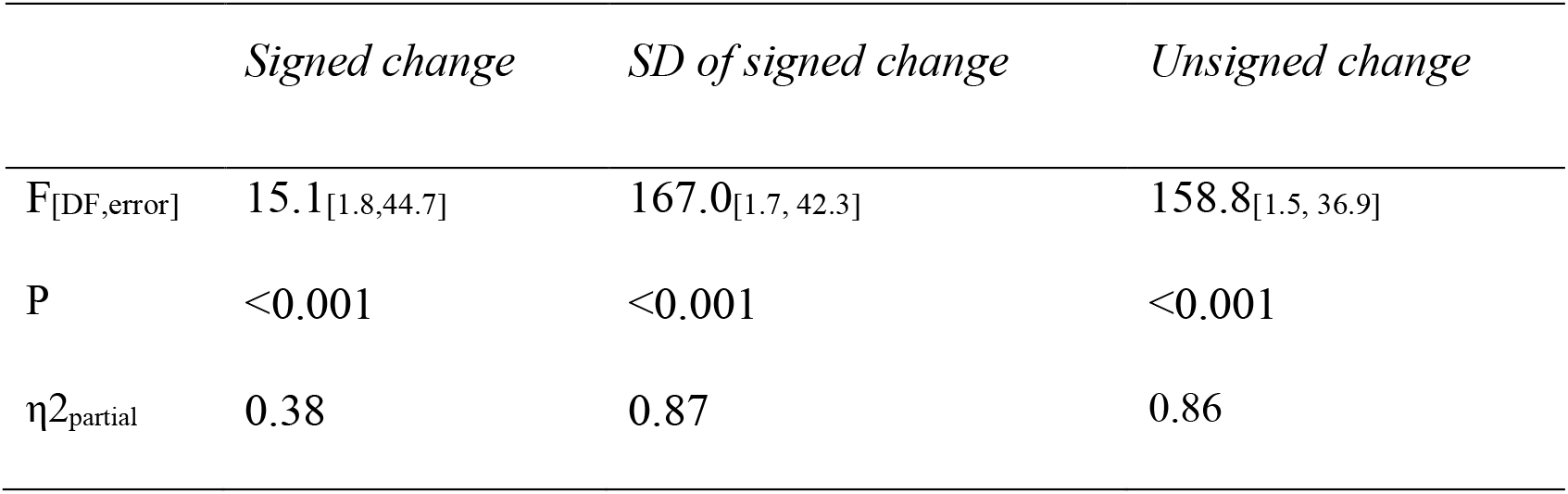
rmANOVA results for the effect of the four outcome histories on the different dependent variables

Post hoc tests were consequently used. For this purpose, we compared the TTV in movement endpoint between S-^(n)^ & S+^(n-1)^ and S-^(n)^ & S-^(n-1)^ and between S+^(n)^ & S+^(n-1)^ and S+^(n)^ & S-^(n-1)^ trials.

The signed TTV (S+^(n)^ & S+^(n-1)^: −0.3° ± 0.5°; S+^(n)^ & S-^(n-1)^: −0.6° ± 1.0°), SD of the TV (S+^(n)^ & S+^(n-1)^: 4.4° ± 1.1°; S+^(n)^ & S-^(n-1)^: 5.0° ± 1.3°) and unsigned TTV (S+^(n)^ & S+^(n-1)^: 3.3° ± 0.8°; S+^(n)^ & S-^(n-1)^: 3.9° ± 0.8°) were significantly different between S+^(n)^ movements that were preceded by S+^(n-1)^ and those that were preceded by S-^(n-1)^ (paired t-test: signed TTV: t = −1.6, P = 0.112; F-test: SD of TTV: F = 13.7, P = 0.001; F-test: unsigned TTV: F = 36.3, P < 0.001, Fig. 3e). Likewise, there were significant differences in signed TTV (S-^(n)^ & S+^(n-1)^: 1.3° ± 1.6°; S-^(n)^ & S-^(n-1)^: −0.1° ± 0.6°), SD of TTV (S-^(n)^ & S+^(n-1)^: 8.9° ± 2.4°; S-^(n)^ & S-^(n-1)^: 14.9° ± 2.9°) and unsigned TTV (S-^(n)^& S+^(n-1)^: 7.0° ± 1.7°; S-^(n)^ & S-^(n-1)^: 11.9° ± 2.7°) after S- movements ^(n)^ between trials that were conditioned on S+^(n-1)^ and S-^(n-1)^ (paired t-test: signed TTV: t = −3.8, P = 0.001; F-test: SD of TTV: F = 96.1, P < 0.001; F-test: unsigned TTV: F = 84.6, P < 0.001, Fig. 3f). These results demonstrate that S+ and S- outcomes in trial^(n-1)^ have differential influences on changes in movement end point after S+ and S- outcomes in trial^(n)^.

We tested whether the differences in TTV conditioned on the past two trials were different for S+^(n)^ and S-^(n)^ trials. Indeed, differences in TTV between S+^(n)^ and S-^(n)^ movements were greater when the previous outcome was S-^(n)^ than when it was S+^(n)^ (paired t-test: TTV: t = −2.4, P = 0.022; F-test: SD of TTV: F = 63.9, P < 0.001; F-test: unsigned TTV: F = 64.9, P < 0.001). These results suggest different RL signals after distinct outcome histories. When participants experienced S+^(n)^ movements, the outcome of the second-to-last trial^(n-1)^ became less influential. Additionally, in case of S-^(n)^ movements, the experience of S- outcomes in trial^(n-1)^ lead to greater motor exploration but to less motor exploration when trial^(n-1)^ was S+.

### PFC oscillatory responses to outcomes during reinforcement motor learning

Since TTV clearly depends on the different outcomes (as evident from the behavioural results), it is indeed plausible that the brain generates different reinforcement signals accordingly to regulate future behavioural adjustments during reinforcement motor learning. Here, we tested this assumption and analysed modulations in neural oscillations during outcome processing.

We concentrated on pre-selected frequency ranges (theta: 4 – 8 Hz, high beta: 25 – 35 Hz) and the time of outcome processing (250 ms – 550 ms after outcome information). First, we plotted the power data in sensor space and observed the greatest increases over frontal sensors for both frequencies (theta: Fig 4a, high beta: 4b). Subsequent source space analyses confirmed our initial assumption that the greatest power increases (relative to pre-feedback) were observed over the prefrontal cortex (theta: Fig 4c, high beta: 4d). In the remaining analyses, we focused on oscillatory activity in the superior frontal gyrus (SFG) and rostral middle frontal gyrus (RMFG), two areas that were previously shown to engage in RL (Garrison *et al*., 2013) (Supplementary Fig. 3). Finally, we extracted averaged time frequency data of the ROI and found similar power time courses as in the sensor space (theta: Fig 4e, high beta: 4f). The data support the assumption that information about motor outcomes result in changes neural oscillations in the PFC.

**Figure 4.**
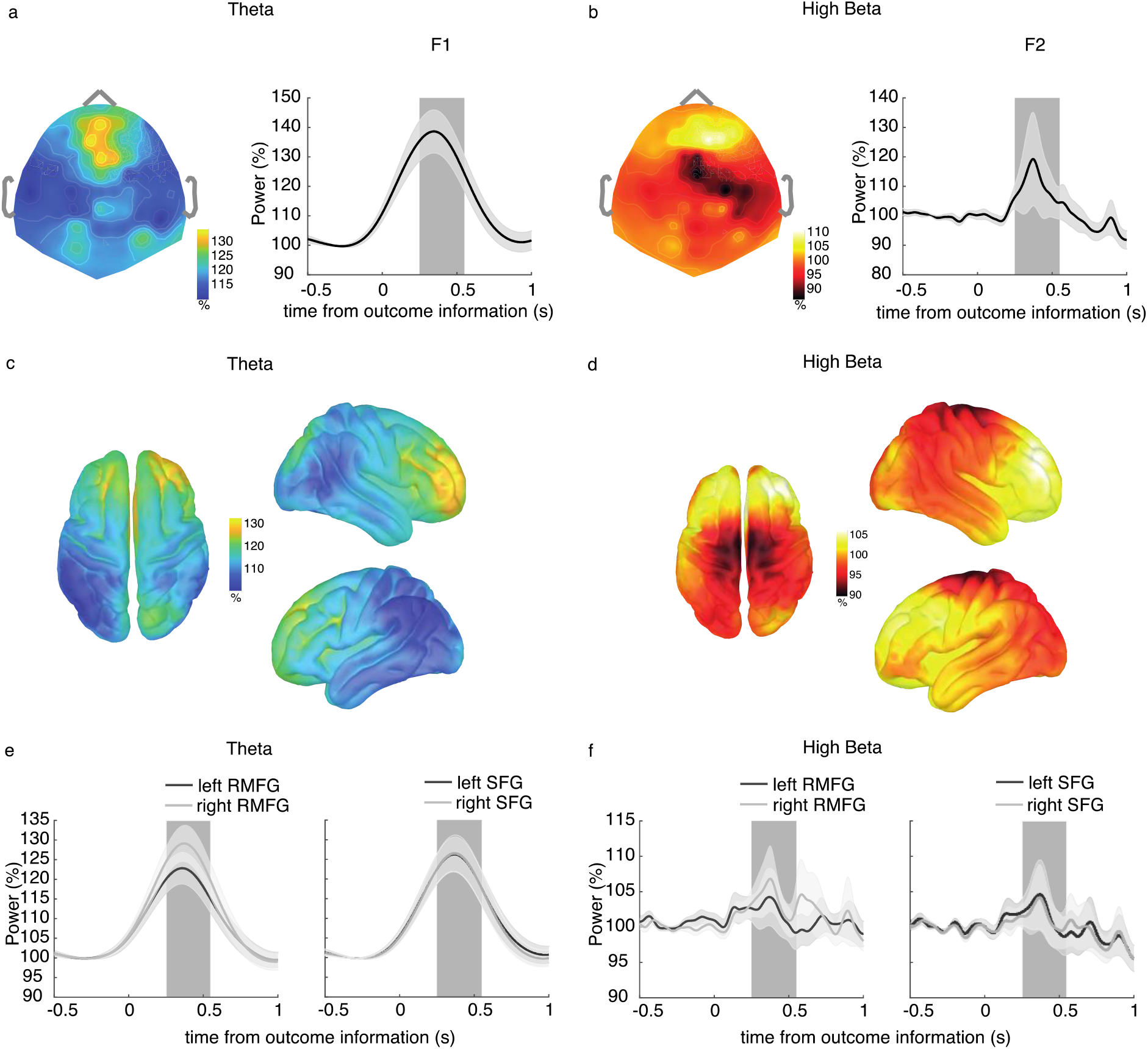
Oscillatory responses in PFC to outcome presentation during reinforcement motor learning. **a & b** Topographical distribution of theta (a) and high beta (b) power during outcome processing (250 – 550 ms after outcome information) (left panel). Grand average power time courses for theta (a) and high beta (b) frequencies during outcome processing (−0.5 s – 1 s relative to outcome information) (right panel). The data show the mean of all trials independent of the prior outcome. Data were plotted for the channel displaying the greatest power increase relative to pre-feedback (theta: F1, high beta: F2). Note that we used different colours to plot the topographical power distribution for theta and high beta frequencies since the data have different scales. **c & d** Source localisation results of theta power (c) and high beta power (d) during outcome processing (average from 250 – 550 ms after outcome information). Data were interpolated on the MRI template. Data are shown from the left, right and front. **e & f** Power time courses of the ROI during outcome processing (−0.5 s – 1 s relative to outcome information). Theta power (e) time courses of bilateral RMFG are shown in left panel and of the bilateral SFG in the right panel. High beta (f) power time courses of bilateral RMFG are shown in the left panel and of the bilateral SFG in the right panel. The shaded rectangle highlights the time window of interest 250 ms – 550 ms. Shading around the mean represent the standard error of the mean.

### Different oscillatory reinforcement signals following S+^(n)^ and S-^(n)^ movements

If oscillatory responses in SFG and RMFG in response to the outcome represent different reinforcement signals, then we would expect different oscillatory responses to S+ and S- outcomes in trial^(n)^. The time courses of theta and beta power changes after both outcomes are shown in Fig 5. Presentation of movement outcomes had differential effects on theta and high beta frequencies depending on the outcome of the movement.

**Figure 5.**
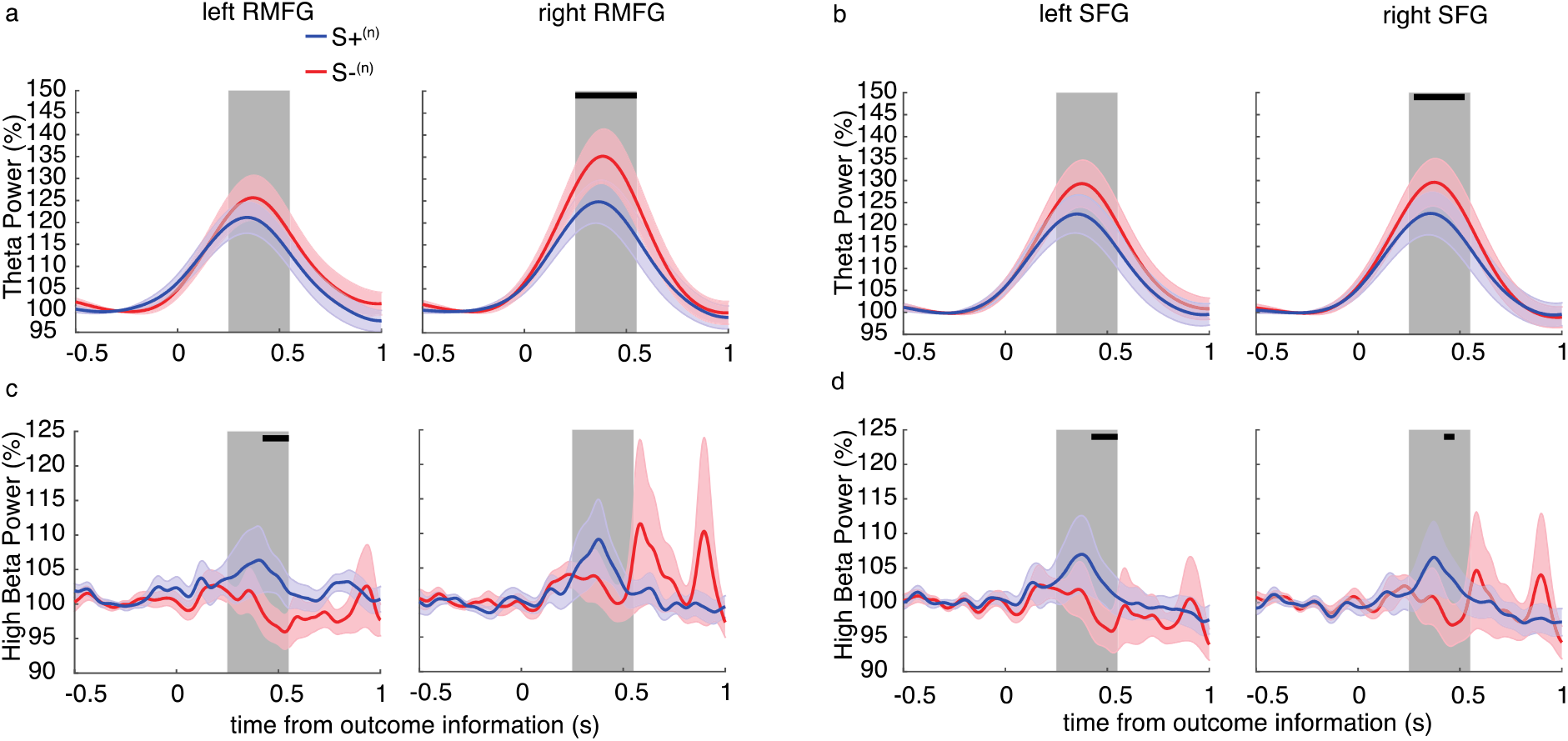
S+(n) and S-(n) outcomes evoke different oscillatory signals in PFC. **a** Theta power time courses of the left RMFG (left panel) and right RMFG (right panel) during outcome processing for S+^(n)^ and S-^(n)^ movements. **b** Theta power time courses of the left SFG (left panel) and right SFG (right panel) during outcome processing for S+^(n)^ and S-^(n)^ movements. **c** High beta power time courses of the left RMFG (left panel) and right RMFG (right panel) during outcome processing for S+^(n)^ and S-^(n)^ movements. **d** High beta power time courses of the left SFG (left panel) and right SFG (right panel) during outcome processing for S+^(n)^ and S-^(n)^ movements. In all plots. the shaded rectangle highlights the time window of interest 250 ms – 550 ms. Shading around the mean represent the standard error of the mean. Significant differences between the outcomes is indicated by the horizontal lines.

In comparison to S+^(n)^ movements, S-^(n)^ movements resulted in greater theta oscillations. While there were no significant differences in theta power over the left RMFG (Fig. 5a, left panel) and left SFG (Fig. 5b, left panel) (all P > 0.05), significant differences in power between S+^(n)^ and S-^(n)^ movements were revealed between 250 ms and 550 ms in the right RMFG (critical P-value: 0.037, Fig. 5a, right panel) and between 300 ms and 500 ms in the right SFG (critical P-value: 0.034, Fig. 5b, right panel). In contrast, S+ movements^(n)^ resulted in greater high beta oscillatory responses compared to S-^(n)^ movements. This was the case in the left RMFG (critical P-value: 0.009, Fig. 5c, left panel) and left SFG from 450 ms and 550 ms (critical P-value: 0.015, Fig. 5d, left panel) and in the right SFG at 450 ms (critical P-value: 0.002, Fig. 5d, right panel). No significant differences in high beta oscillatory response between S+^(n)^ and S-^(n)^ movements were observed in the right RMFG (P > 0.05) (Fig. 5c, right panel). Similar results were obtained for sensor-space data (Supplementary Fig. 4). The data suggests different oscillatory responses to S+^(n)^ compared to S-^(n)^ outcomes in the prefrontal cortex. While S+^(n)^ outcomes lead to greater high beta band power, S-^(n)^ outcomes lead to greater theta band power.

### Oscillatory reinforcement signals in PFC depend on the outcomes of the past two movements

Next, we asked whether neural oscillations during outcome processing depend on the outcome of the past two trials. Due to differences in the number of trials per each outcome scenario and potential differences in signal-to-noise ratios, we performed a bootstrapping with replacement for each individual EEG dataset. The bootstrapped grand average time frequency responses of our ROI are shown in Fig. 6.

**Figure 6.**
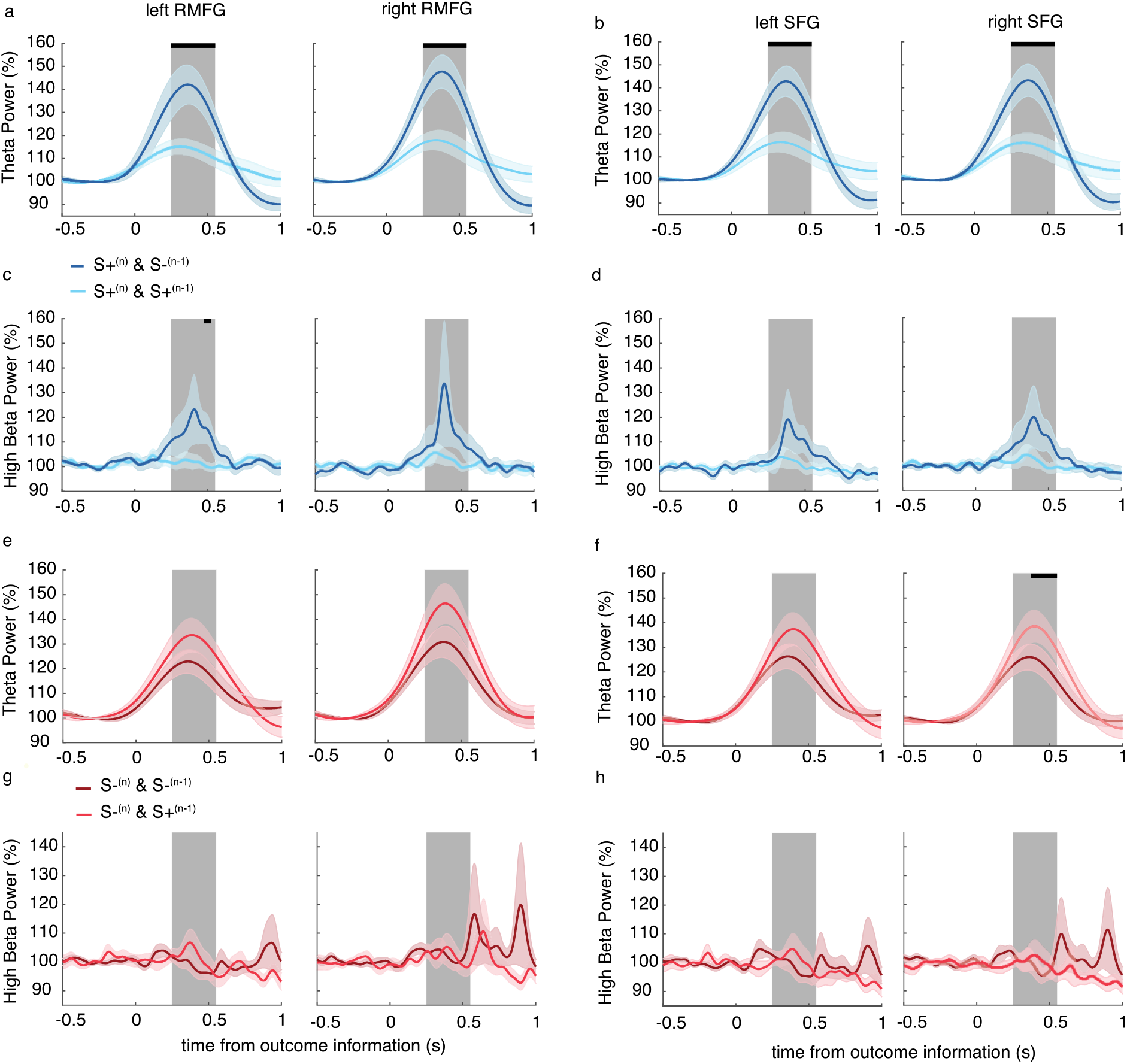
Oscillatory reinforcement signals in PFC depend on the outcome history. **a** Theta power time courses of the left RMFG (left panel) and right RMFG (right panel) during outcome processing for S+^(n)^ movements that were preceded by S+^(n-1)^ movements and S-^(n-1)^ movements. **b** Theta power time courses of the left SFG (left panel) and right SFG (right panel) during outcome processing for S+^(n)^ movements that were preceded by S+^(n-1)^ movements and S-^(n-1)^ movements. **c** High beta power time courses of the left RMFG (left panel) and right RMFG (right panel) during outcome processing for S+^(n)^ movements that were preceded by S+^(n-1)^ movements and S-^(n-1)^ movements. **d** High beta power time courses of the left SFG (left panel) and right SFG (right panel) during outcome processing for S+^(n)^ movements that were preceded by S+^(n-1)^ movements and S-^(n-1)^ movements. **e** Theta power time courses of the left RMFG (left panel) and right RMFG (right panel) during outcome processing for S-^(n)^ movements that were preceded by S+^(n-1)^ movements and S-^(n-1)^ movements. **f** Theta power time courses of the left SFG (left panel) and right SFG (right panel) during outcome processing for S-^(n)^ movements that were preceded by S+^(n-1)^ movements and S-^(n-1)^ movements. **g** High beta power time courses of the left RMFG (left panel) and right RMFG (right panel) during outcome processing for S-^(n)^ movements that were preceded by S+^(n-1)^ movements and S-^(n-1)^ movements. **h** High beta power time courses of the left SFG (left panel) and right SFG (right panel) during outcome processing for S-^(n)^ movements that were preceded by S+^(n-1)^ movements and S-^(n-1)^ movements. In all plots. the shaded rectangle highlights the time window of interest (250 ms – 550 ms after outcome information). Shading around the mean represent the standard error of the mean. Significant differences between the outcomes is indicated by the horizontal lines.

Theta power was significantly greater between 250 ms and 550 ms in the left RMFG (critical P-value: 0.005, Fig. 6a, left panel), the right RMFG (critical P-value: 0.001, Fig. 6a, right panel), the left SFG (critical P-value: 0.003, Fig. 6b, left panel) and the right SFG (critical P-value: 0.001, Fig. 6b, right panel) in S+^(n)^ movements that were preceded by S-^(n-1)^ movements than those that were preceded by S+^(n-1)^ movements. Likewise, high beta power was significantly greater at 500 ms in the left RMFG (critical P-value: 0.005, Fig. 6c, left panel) in S+^(n)^ movements that were preceded by S-^(n-1)^ movements than those that were preceded by S+^(n-1)^ movements. Though there were similar tendencies in the other regions, the results did not reach statistical significance (P > 0.05) in the right RMFG (Fig. 6c, right panel), the left SFG (Fig. 6d, left panel) and the right SFG (Fig. 6d, right panel).

Finally, the time course of theta power was not significantly different between S-^(n)^ movements that were preceded by S-^(n-1)^ movements and those that were preceded by S+^(n-1)^ movements in the left RMFG (Fig. 6e, left panel), the right RMFG (Fig. 6e, right panel) and the left SFG (Fig. 6f, left panel) (all P > 0.05). Theta power between 400 ms and 550 ms was significantly greater in S-^(n)^ movements that were preceded by S+^(n-1)^ movements compared to those that were preceded by S+^(n-1)^ movements in the right SFG (critical P-value: 0.028, Fig. 6f, right panel). No significant differences were observed for high beta power in the left RMFG (Fig. 6g, left panel), the right RMFG (Fig. 6g, right panel), the left SFG (Fig. 6h, left panel) and the right SFG (Fig. 6h, right panel) between S-^(n)^ movements that were preceded by S+^(n-1)^ movements and those that were preceded by S+^(n-1)^ movements (all P > 0.05). Additional information on sensor-space data can be found in Supplementary Fig. 5. The results suggest that high beta band power increases especially in cases were S+^(n)^ trials are preceded by S-^(n-1)^ trials. On the contrary, theta band power increases were most prominent when the outcomes in two subsequent trials changed (from S+^(n-1)^ to S-^(n)^ or from S-^(n-1)^ to S+^(n)^).

## Discussion

The present study revealed outcome-specific adjustments of TTV in movement endpoint during reinforcement motor learning. We found that S+^(n)^ movements were followed by relatively little TTV and a greater proportion of S+ movements in the subsequent trial^(n+1)^. In contrast, S-^(n)^ movements caused increased TTV and had a lower proportion of S+ movements in the subsequent trial^(n+1)^. We investigated the regulation of TTV in further detail and found that TTV following S+^(n)^ movements was less influenced by the outcome of the previous trial^(n-1)^. In contrast, TTV was greater after S-^(n)^ movements when they were preceded by S-^(n-1)^ trials compared to when they were preceded by S+^(n-1)^ trials. These results suggest a change in values of previous outcomes when human participants experience S+^(n)^ and S-^(n)^ movements during a reinforcement-based motor learning task. We hypothesised that these behavioural effects involve different cortical reinforcement signals and analysed neural oscillations from the prefrontal cortex (bilateral RMFG and SFG). In general, S+^(n)^ movements led to increased high beta oscillatory activity while S-^(n)^ movements caused greater theta oscillatory activity. However, when the power data were conditioned on the past two trials we found that these oscillatory differences are driven by distinct outcome histories. Beta power following feedback presentation was greatest in S+^(n)^ movements that were preceded by S-^(n)^ movements providing a potential mechanism to reinforce the current S+^(n)^ movement and disregard the behaviour from the previous S-^(n-1)^ trial. Theta oscillatory activity was greatest after S-^(n)^ movements when the preceding movement was S+^(n-1)^ and after S+^(n)^ movements when the preceding movement was S-^(n-1)^ suggesting that increased theta oscillations reflect error detection, i.e. large discrepancies between expected and actual outcomes (reward prediction errors) and thereby a change in the variability state. Therefore, our results provide evidence for an outcome-specific regulation of TTV and oscillatory reinforcement signals in the prefrontal cortex during human reinforcement motor learning.

Learning from the outcome of recent trials is an important component of motor learning and helps to adjust future motor behaviour. For instance, reinforcement of S+ movements through rewards causes stable performance gains (Pekny *et al*., 2011) and results in greater motor retention than learning from punishment (Galea *et al*., 2015). Negative (punishment) feedback helps us to avoid S- movements and accelerates online improvements in motor performance (Galea *et al*., 2015). These results suggest that learning from S+ and S- outcomes involves distinct processes. In the present study, we showed that S+ and S- motor outcomes lead to different adjustments in TTV. In agreement with an earlier report (Pekny *et al*., 2015), TTV was decreased after S+ movements and increased after S- movements. The outcome-dependent modulation of TTV was additionally an important aspect of efficient reinforcement motor learning as individual differences in TTV (after S+^(n)^ trials but not after S-^(n)^ trials) predicted overall motor performance.

However, it is unlikely that updating of motor behaviour uses only the outcome of the current action. Rather, motor variability is causally regulated by recent trial outcomes in rats, that is the integrated outcomes of the past ∼10 trials (Dhawale *et al*., 2019). The relevance of trial outcome on motor variability decays in an exponential weighted manner, i.e. very recent outcomes have a greater effect compared to “older” outcomes. In the present study, we focused on the effects of the past two trial outcomes.

In cognitive decision-making tasks, participants are often presented a finite number of available options, e.g. in gambling games (Cohen *et al*., 2007; HajiHosseini & Holroyd, 2015). Since there is no direct association between the values of the different options, choosing an option does not inform about the value of the other options in these tasks. This is different in the motor domain as the agents act in a continuous motor space (Dhawale *et al*., 2019). Here, a S+ movement end point implies that neighbouring movement end points likely have a similar positive value. In contrast, an S- movement suggests that S+ movement end points are located at more distant positions. However, the value of previous outcomes may depend on the acutely experienced outcomes. Here, we showed that TTV is less influenced by the outcome of the second-to-last trial when the current trial was S+. The findings suggest that previous outcomes are assigned less value as soon as humans experience positive motor outcomes, a process that might promote acute improvements in short-term motor performance. In contrast, TTV was increased after S- movements, especially when the second-to-last trial was also S-. S- trials that were preceded by S+ trials showed less TTV. This implies that participants consider previous S+ outcomes for adjustments in TTV, even when they acutely experience S- outcomes. We speculate that motor outcomes prior to positive outcomes are disregarded since they contain only little relevant information on potential future behavioural adjustments in the task.

The efficacy of this mechanisms may depend on the certainty of the task and environment (Dhawale *et al*., 2019). In task situations where the conditions are stationary, a greater reliance on previous movements might be a reasonable strategy to reproduce the positive outcomes. In non-stationary conditions, too much reliance on previous positive outcomes may be ineffective since the task conditions continuously change and new task solution have to be explored.

Our results imply that S+ outcomes reinforce recent behaviour and inform an exploitation strategy (low variability state) while S- outcomes will stimulate and inform an exploration strategy (high variability state). These distinct motor states might be differently encoded at the level of neural oscillations. Here, we demonstrate that prefrontal neural oscillations in theta and high beta frequencies respond selectively to different outcome histories during reinforcement motor learning, a mechanism that potentially gates future adjustments in motor output. Activity in the prefrontal cortex is sensitive to distinct motor states, characterized by marked differences in variability. For instance, activity in regions of the PFC is modulated when participants switch between exploratory and exploitative behavioural modes during decision-making tasks (Daw *et al*., 2006). Moreover, neural activity in the prefrontal cortex is sensitive to the recent history of rewards, which might function as an update on estimates of predicted rewards (Kim & Shadlen, 1999; Barraclough *et al*., 2004; Padoa-Schioppa & Assad, 2006; Rushworth & Behrens, 2008; Seo & Lee, 2008; Histed *et al*., 2009). The cerebral regions that are involved in RL include prefrontal cortical areas such as SFG, RMFG and cingulate cortex (Garrison *et al*., 2013). Recently, it has been suggested that the prefrontal cortex may serve as a meta-reinforcement learning system, which is controlled by the midbrain dopamine system but acts as an independent learning system (Wang *et al*., 2018). Previous human decision-making studies (involving e.g. gambling tasks) have shown that neural oscillations in prefrontal cortex respond to reward-related feedback stimuli (Luft, 2014). Neural oscillations reflect the synchronous activity of neural assemblies and have been considered important to integrate large-scale networks (Fries, 2005), to facilitate synaptic plasticity during learning (Buzsaki & Draguhn, 2004) and to engage in the control of top-down information flow (Engel *et al*., 2001). These studies suggest that the PFC might also be Involved in processing the outcome of previous motor actions that can be used to guide subsequent movements during motor tasks.

The results of the present study support this assumption by showing that neural oscillations in the SFG and RMFG respond selectively to positive and negative motor experiences. We showed that theta oscillations were most prominent after S- movements when they were preceded by S+ outcomes and after S+ movements when they were preceded by S- movements. Theta oscillations are present during cognitive tasks and modulated by attentional and working memory demands (Kubota *et al*., 2001; Onton *et al*., 2005). The increase could reflect an increased cognitive load that is caused by the detection of conflicts and/or errors, e.g. a mismatch between predicted and actual rewards and thereby induce a change in the variability state (from low TTV to high TTV or vice versa).

In contrast, high beta oscillations in prefrontal cortex were most prominent after S+ movements; and especially so when they were preceded by an S- movement. This indicates that the reinforcement signal is indeed more distinct when the learner experiences unexpected (positive) outcomes (Akitsuki *et al*., 2003; HajiHosseini *et al*., 2012). The increased beta activity could constitute a potential mechanism that ensures that the current S+ action is reinforced while previous negative actions are assigned less value and, in this way, beta activity could inform the model of future motor behaviour. In other words, greater beta oscillations in the prefrontal cortex after S+ trials reinforce the most recent motor behaviour and inherently also signal less weight on previous outcomes in planning of the next movement.

It is likely that oscillations in different frequency ranges and neuronal circuits interact to support a common goal. For example, the interaction of theta and beta band oscillations has been proposed to process and store short-term memories (Lisman & Idiart, 1995; Axmacher *et al*., 2010). Likewise, increases in the functional connectivity between the striatum and the prefrontal cortex have been observed during categorical learning (Antzoulatos & Miller, 2014). A role for the dopaminergic system is supported by the fact that activity of dopaminergic neurons is modulated by reward stimuli in primates (Mirenowicz & Schultz, 1994). Future studies have to clarify how interactions between different neural circuits engage in RL.

The present study contains a number of limitations that should be considered. Due to the experimental requirements when combining reinforcement motor learning with EEG measurements in the present study, we were not able to sample more than 320 movements per participant in the main protocol. Based on this, it was not feasible to analyse further outcome sequences including “older” outcomes since these would have contained a considerably smaller number of trials. However, on the basis of a previous study in rats (Dhawale *et al*., 2019), it can be assumed that “older” outcomes also have an impact on the behavioural trial-to-trial adjustments during reinforcement motor learning in humans.

Based on the behavioural measures, we are not able to discern to which extent changes in TTV are the consequence of intended motor variability or simply a by- product of sensorimotor noise or error in motor acuity i.e. unintended motor variability. While changes in TTV after S- trials can be due to sensorimotor noise and active exploration, changes in TTV after S+ trials should be mainly caused by sensorimotor noise (van Mastrigt *et al*., 2020). Thus, motor variability is influenced by a number of factors including intended and unintended variability regulatory mechanisms (Pekny *et al*., 2015; Therrien *et al*., 2018; Dhawale *et al*., 2019; van Mastrigt *et al*., 2020). Nevertheless, the results support the notion that negative outcomes inform an explorative strategy leading to higher TTV whereas positive outcomes lead to exploitation, which is characterized by low TTV.

We focused our analyses of neural oscillations to prefrontal cortical regions since we had a strong a-priori hypothesis that a prefrontal cortical network is involved in RL. It is however very likely that the cortical activity is influenced also by other circuits involving the basal ganglia, which also play an important role in reward-based learning. For instance, regulation of motor variability is impaired after inactivation of a cortico-basal ganglia circuit of songbirds (Olveczky *et al*., 2005) and in human Parkinson patients (Pekny *et al*., 2015). Future studies should investigate the interactions of the different network nodes and their influence on motor output. While the present results do not provide a causal link between the behavioural observations and the modulations in neural oscillations, previous studies do however suggest a clear association between modulations in neural oscillations and behavioural adjustments (Luft, 2014). In conclusion, we provide evidence that positive outcomes “overwrite” previous motor states to a greater extent than negative outcomes and that changes in high beta and theta oscillatory activity in the prefrontal cortex potentially reflect changes in the movement variability state during reinforcement motor learning. Therefore, the results provide novel insights on the behavioural and neural mechanisms that underline learning from previous outcomes.

## Methods

### Participants and study design

Twenty-six participants were recruited for the study by convenience sampling using flyers and social media. All participants were healthy young adults (mean age: 25.4 ± 2.7 years old, 16 females). Participants had to be free of any neurological or psychiatric diseases and have normal or corrected-to normal vision to be included in the study. According to the Edinburgh Handedness questionnaire (Oldfield, 1971), participants were right-handed (laterality index of 82.2 ± 42.4). The study was approved by the regional ethical committee (H-17019671). The study conformed to the standards set by the Declaration of Helsinki (latest revision in Fortaleza, Brazil). All participants gave their written informed consent to the procedures of the study prior to participation. In summary, each participant performed a reinforcement motor task and experienced different outcomes (target hit or miss) dependent on their motor performance. We compared the effects of the outcome on behavioural and oscillatory responses in an experimental within-participant design with random effects.

### Experimental setup

Participants sat in a laboratory chair approximately 50 cm in front of a computer screen (27” monitor with 60 Hz frame rate and 2,560 x 1,440-pixel resolution). The height of the computer screen was individually adjusted. Participants positioned their left forearm in a neutral position in a splint that was placed on a table next to them. The left hand grabbed a handle with a built-in goniometer and could be moved by performing wrist flexion/extension movements. The forearm and hand position were stabilized and supported with Velcro® straps to avoid changes in elbow and shoulder joint angles. The view of the forearm and hand was hidden by a custom-built box to prevent visual feedback of the moving hand. Goniometer data were recorded at a sampling frequency of 2048 Hz (CED 1401+ with Signal 3.09 software, Cambridge Electronic Design Ltd., UK) and stored offline for further analyses.

### Motor task

All participants performed wrist flexion movements with their left wrist (Fig. 1a). The goal of the task was to move a circular cursor (radius of 15 pixels) into a target area on a computer screen. The position of the target area was changed several times during the experiment. There were three different target positions (see experimental protocol) and each target had a horizontal size of 330 pixels (green target: 1130 to 1460 pixels, grey target: 1680 to 2010 pixels, purple target: 2230 to 2560 pixels) (Fig. 1b).

In each trial, participants performed one wrist flexion movement. Visual traffic lights and text on the computer screen marked the beginning of a trial (Fig. 1c). The computer cursor always started on the left side of the screen. The start of a trial was signalled by the appearance of a red dot that was positioned on the left side of the screen. After 500 ms a yellow dot appeared underneath the red dot and instructed participants to prepare the movements. After another 500 ms a green dot appeared as the final GO cue. Next, participants could move the cursor to the right by performing wrist flexion movements.

Participants were instructed to not adjust the movement end position once they finished their movement. To avoid post-movement corrections, cursor motion was insensitive to wrist movements once the differences in cursor position fell below five pixels in five consecutive samples. Since the goal of the motor task was an accurate movement end position, participants were informed prior to the experiment that the task was not a reaction time task. Binary feedback about motor performance (S+ or S-) was given 2500 ms after the GO signal for 1000 ms. Participants were explicitly instructed to relax their arm during that period. After information of the outcome, participants were instructed to move back to the starting position and wait for the start of the next trial. If flexion movement time was longer than 800 ms in two consecutive trials, we asked participants to move faster in the next trial. The next trial was started after 5500 ms. Thus, each trial lasted 10 s.

A movement was considered S+ if the centre of the cursor was within the target area. Success was indicated by a text box indicating “Good job”. After S- trials, participants were prompted with a text box stating “Try again”. When the trial was S+, a point was added to the participant’s score. The participant’s score was displayed continuously throughout a training block (40 trials) with the aim of motivating the participants in each block. All participants were compensated equally for the time they had spent in the laboratory and compensation was not based on their motor performance. The positions and size of the target areas were determined in pilot experiments (n = 8, results are not reported in this paper) such that participants yielded on average approximately an equal number of S+ and S- trials. The task was created using MATLAB R2019a (The MathWorks, Natick, Massachusetts, United States, R2019a) and the Psychophysics Toolbox extensions (Brainard, 1997). Cursor movements (in pixels), distance from the targets (in pixels) and information on movement outcome for each trial were logged online and saved for further offline analyses.

### Experimental protocol

Prior to the study, all participants were informed about the experimental protocol and the motor task. Subsequently, participants were introduced to the task by performing ten wrist flexions with no targets displayed on the screen. After this short introduction, participants performed 40 movements with online visual feedback (Fam1) and 40 without online visual feedback (Fam2) of cursor motion and target position. In both familiarization blocks, participants had to reach for the grey middle target and they received offline binary feedback about their performance at the end of the trial (knowledge of result). The target position did not change during these blocks. These initial blocks were performed to ensure that participants could adhere to the time course of a trial, to familiarize them with the handle’s sensitivity and to quantify baseline movement variability.

In the main protocol, participants performed 320 wrist flexion movements in blocks of 40 movements. Between blocks, we incorporated breaks of 2 minutes to ensure that participants focused on the motor task the whole experiment. Pilot experiments demonstrated that for most participants 40 consecutive movements (approx. 7 minutes) was feasible and could be performed with sufficient and sustained attention. Moreover, the target position was changed several times during the experiment. The green and purple targets were present in 80 trials, respectively, (i.e. 160 trials) and the grey target in the remaining 160 trials. Thus, in total, participants performed 160 movements to an already familiarized target position (grey target) and 160 movements to an unfamiliar target position (green or purple target). Targets were changed every 25, 30 or 40 trials such that there was no systematic order of the different conditions. The size of the target area remained constant throughout the experiment.

In the main protocol, participants did receive knowledge of result but not knowledge of performance. All participants were informed that the target position could change at any time during the experiment. Giving this information, we wanted to avoid too much frustration (which could influence participant’s motivation) but also stimulate the exploration of different motor actions following a history of unsuccessful trials.

### Pre-processing of behavioural data

In the offline analysis, we analysed movement kinematics from the goniometer signal of the wrist. In brief, continuous data was epoched from −1000 ms to 3000 ms relative to the GO cue to capture start and end points of all movements. For these epochs, we calculated the first derivate of the goniometer signal for all single trials. Subsequently, we smoothed the derivates by calculating the moving average of 400 samples and applying a 3^rd^ order Butterworth lowpass filter (20 Hz). We used the smoothed signals to determine the movement start and end points by calculating the maximum movement speed within each epoch. Movement start was defined as the sample for which signals exceeded 15% of its respective speed maximum. Movement end was defined as the sample for which the signals dropped below 15% of its respective speed maximum. The movement end point angle was calculated by subtracting movement end angles from movement start angles (in °). During this process, all data were continuously visually inspected and checked. In addition to the kinematic analysis, we also extracted the information on trial outcomes (hits or misses). Trials in which the participants did not move at all were discarded (movement angle < 2°).

### Behavioural data analyses

First, we were interested in the association of the outcome time series (S+ and S- trials) with lagged versions of the same outcome time series. Thus, we performed partial autocorrelation (PAC) for each participant (Matlab function: *parcorr*). This analysis was performed to obtain an initial estimate of the impact of previous outcomes on future outcomes and to test the assumption that the past two outcomes have the greatest association with future outcomes.

In the remainder of our analysis, we focused on changes in movement endpoint (Δμ) as a function of outcomes from the previous two trials. There were two reasons for restricting the analyses to the last two outcomes. First, the PAC analysis confirmed that the past two outcomes have the greatest correlation with future outcomes. Second, outcome histories considering even “older” actions would have contained too few trials (<10) to perform valid comparisons.

In the following, trial^(n)^ refers to the most recent trial and trial^(n+1)^ to the subsequent trial. We measured the differences in movement endpoint (Δμ) between trial^(n+1)^ and trial^(n)^ as a measure of TTV using the following formula:

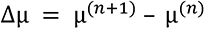

For each participant, we extracted the signed TTV in movement endpoint (Δμ) conditioned on the outcome of the previous trial^(n)^. The signed TTV gives an estimate on the directional changes in movement endpoint and potentially reveal whether changes in movement endpoint are biased in one or the other direction. To quantify unsigned TTV in movement endpoint, we also calculated the absolute TTV in movement endpoint (|Δμ|). The unsigned TTV is a measure of the absolute motor exploration independent of the direction of the change.

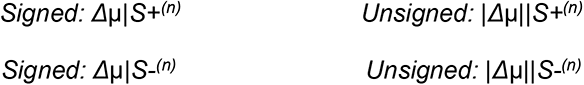

The first movement of every block were discarded, as these had no previous trial. Subsequently, individual data histograms from the different conditions were visually inspected. In accordance with a previous study (Pekny *et al*., 2015), signed data showed a normal distribution and unsigned data had a negative exponential distribution. Thus, we fitted normal distributions as the conditional probability distribution for the signed data p(Δμ|S+^(n)^) and p(Δμ|S-^(n)^). For the unsigned data, we fitted exponential distributions as the conditional probability distribution p(|Δμ||S+^(n)^) and p(|Δμ||S-^(n)^). The mean (M) and standard deviation (SD) were calculated from the individual normal distributions as a measure of the signed TTV and the variations in the TTV, respectively. The M was calculated from the individual exponential distribution as a measure of the absolute TTV.

In the next step, we asked whether TTV is dependent on the previous two outcomes, i.e. the most recent trial^(n)^ and the second-to-last trial^(n-1)^. Thus, we extracted TTV from four different outcome histories:

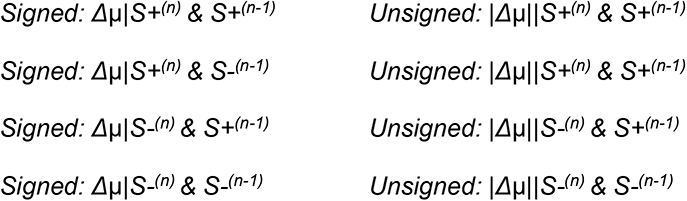

Since these four conditions contained a significantly different number of trials, we performed bootstrapping with replacement for each individual (Matlab function: *datasample*). As suggested previously (Hesterberg, 2011), we sampled 1,000 new datasets per participant and condition, each of them containing 100 randomly chosen trials from the original data. We chose to use this bootstrapping approach since differences in trial numbers can significantly bias measures of variability in the distinct outcome histories (e.g. the SD). From the individual bootstrapped probability distributions, we computed the M and SD as described above to derive robust confidence intervals for estimating standard errors and for hypothesis testing. Again, we fitted normal (Δμ) and exponential (|Δμ|) distributions for changes in movement endpoint after S+*^(n)^* and S-*^(n)^* movements that were either preceded by S+^(n-1)^ or S-^(n-1)^ movements, respectively. Probability distributions were fitted using the MATLAB functions *fitdist* and *makedist.* All described analyses were performed offline using MATLAB R2019a (The MathWorks, Natick, Massachusetts, United States, R2019a).

### Statistical analyses of TTV

Statistical analyses were performed using SPSS software 27 (SPSS®, Chicago, IL, USA) and RStudio (RStudio Team, 2018). All dependent variables were tested for normality by the Kolmogorov-Smirnov test. Homogeneity of variances was tested using the Fisher’s F-test. Repeated-measures analyses of variance (rmANOVA) with random effects were calculated to test the effect of outcomes on the signed and unsigned TTV and the SD of the signed TTV. The rmANOVA design had one within-participant factor with 4 levels defining the outcome history (S+^(n)^ & S+^(n-1)^, S+^(n)^ & S-^(n-1)^, S-^(n)^ & S+^(n-1)^, S-^(n)^ & S-^(n-1)^). The Greenhouse–Geisser correction was used for rmANOVAs if the assumption of sphericity was violated (Mauchly’s test). Effect sizes were estimated using Eta-squared (η2 partial). Within-participant comparisons between the different outcomes were performed with appropriate paired t-tests or F-tests. Paired t-tests were calculated for comparisons of the distance from target (in pixels), the number of S+ and S- movements and the M values from the normal distribution of the signed changes in movement endpoint. F-tests were calculated for of the SD values and the M values from the exponential distribution. The test of equal or given proportions was used to compare the proportion of S+^(n+1)^ movements after the different outcomes. A P-value correction using the Benjamini-Hochberg procedure was applied to control the False discovery rate (FDR). Uncorrected p-values are presented in the Results section. A significance level of P < 0.05 was assumed. All data are reported as M ± SD if not stated otherwise.

### EEG recordings

EEG recordings were performed with a 64 channel Biosemi system (BioSemi, Amsterdam, The Netherlands) using the software ActiView (version 8.06). Electrodes were embedded into an electrode cap and positioned according to the extended 10-20 system. Data recording was performed at 2048 Hz and the online reference was the Common Mode Sense (CMS)/Drive Right Leg (DRL). It was ensured that the electrode offsets (voltage differences between single electrodes and CMS) were less than ±20 μV. EEG signals were recorded continuously throughout the experiment. During the motor task, information of task timing and movement outcome was inputted (via a trigger signals) from Matlab to synchronize the task and the EEG recording and allow time-locked offline analyses of the EEG data. All participants were asked to relax their face and neck muscles during the experiment to minimize signal artefacts.

### EEG pre-processing

EEG data was pre-processed using EEGlab (Delorme & Makeig, 2004). Data were imported and the DC offset was corrected by subtracting the mean from each channel. Noisy channels were identified by visual analysis of the time-series (high amplitude time series, e.g. from tonic muscle activity) and frequency plots of the entire recording period (strong power deviations and/or unusual spikes in the frequency range 0.5 Hz – 48 Hz). Subsequently, we removed 4.48 ± 1.96 channels per participant which were mostly located temporally and/or occipitally. EEG signals were i) re-referenced to the average of all channels, ii) bandpass-filtered between 0.5 Hz and 48 Hz using a FIR filter and iii) down-sampled to 256 Hz. The pre-processed EEG data were segmented into epochs ranging from −3500 ms to +3500 ms relative to the beginning of information of the outcome. Using the segmented data, we performed independent component analyses (ICA) with the runica algorithm. The ICA was performed on segmented data as our continuous recordings also encompassed a great amount of irrelevant data (e.g. breaks). The aim of this step was to identify and remove components indicating horizontal and vertical eye movements and saccades based on the topological and spectral characteristics of the components (Chaumon *et al*., 2015). On average, we selected and removed 2.12 ± 0.33 components per participant. Finally, removed channels were interpolated with the standard spherical method.

All epochs were visually inspected from each participant to identify and remove trials that were contaminated by signal artefacts (e.g. high amplitude signal deviations from strong muscle artefacts in most channels). Following this assessment, we had to remove the data from one participant since more than 25% of all trials had to be excluded. For the remaining 25 participants, we removed on average 4.26 ± 4.4% trials (out of 320) per participant.

For each participant, we created datasets encompassing i) all trials independent of the outcome, ii) S+^(n)^ and S-^(n)^ trials and ii) S+^(n)^ & S+^(n-1)^ trials, S+^(n)^ & S-^(n-1)^ trials, S-^(n)^ & S+^(n-1)^ trials and S-^(n)^ & S-^(n-1)^ trials. All datasets were saved offline for further analyses. The subsequent EEG analyses comprised the following steps. First, we tested the assumption that the information of movement outcome leads to changes in oscillatory activity (normalised to a “pre-feedback” period) in frontal sensors and in prefrontal cortex in source space. Subsequently, we constructed power time courses of two prefrontal cortical regions previously shown to engage in RL, i.e. SFG and rostral middle frontal gyrus RMFG (Garrison *et al*., 2013), to compare power time series between different outcomes. The regions of interest (ROI) are highlighted in Supplementary Fig. 3. In our analyses, we focussed on pre-specified frequency ranges and time windows of interest (see Time-frequency analyses in sensor space).

### Time-frequency analyses in sensor space

Further data analyses were performed on the pre-processed segmented data using Brainstorm (Tadel *et al*., 2019), which is documented and freely available for download online under the GNU general public license (http://neuroimage.usc.edu/brainstorm). All datasets were imported to Brainstorm in the time range −2000 ms to +2000 ms relative to information of the outcome. The reason for this time window was a compromise between avoiding filtering related edge artefacts of time-frequency analyses and minimising computational demands.

All single trials were transformed to the time-frequency domain by convolving the signal with a set of complex Morlet wavelets, which are defined as complex sine waves tapered by a Gaussian. The full-width at half-maximum was 3000 ms and the sine waves were created at 1 Hz (corresponding to 7 cycles). The analyses were constrained to the theta band (4 – 8 Hz) and the higher beta band (25 – 35 Hz). The rationale of this choice was that there is ample evidence that oscillatory modulations in both frequency bands represent different reinforcement signals during learning from feedback (Luft, 2014). For simplicity, the frequency range 25 – 35 Hz will be referred to as high beta although it also includes low gamma frequencies per some definitions.

Single time-frequency series were averaged for each participant and frequency band (theta and high beta) at each electrode. In order to calculate event-related changes in power, the averaged data were scaled with the mean of a “pre-feedback” period prior to outcome information (−400 ms to −200 ms). This period covers the time where participants had already finished the movement and were awaiting outcome information. This normalisation procedure ensures that the data across all time points, sensor/source points, conditions and subjects are in the same scale and hence comparable. Changes in power were calculated in %, that is (x/u) *100 where x is the data and u is the mean over the “pre-feedback” period. This analysis was run on the dataset encompassing all trials independent of previous outcomes. Data were visually inspected by plotting selected sensors and topographical plots on pre-specified time windows (250 – 550 ms) on the basis of previous studies (Cohen *et al*., 2007; Marco-Pallares *et al*., 2008; HajiHosseini *et al*., 2012; Luft, 2014; HajiHosseini & Holroyd, 2015; Marco-Pallares *et al*., 2015).

### Source modelling

Motivated by the results in sensor-space, we performed source analyses on the pre-processed sensor time series to localize and reconstruct the cortical regions contributing to oscillatory modulations during RL of motor skills. In a first step, we created our forward model describing how neuronal activity propagates from each cortical position to the EEG sensors, also called the lead field matrices. Since individual MRIs were not available, we used the MNI International Consortium of Brain Mapping152 brain template which is a non-linear average of 152 participants (Fonov *et al*., 2009). The forward model was constructed using the symmetric boundary element method (BEM) from the open-source software OpenMEEG (Gramfort *et al*., 2010). The BEM uses three realistic layers (head, outer skull and inner skull; 1922 vertices per layer) to calculate the volume conduction model. The relative conductivity of the layers was [1 0.0125 1] which describes the relative conductivities of each layer and is the default in Brainstorm. Standard BioSemi sensor positions were aligned to the template head space. The source space contained 15002 elements constrained to the cortical sheet. The number of vertices has been suggested to be sufficient to sample the folded details of the cortex (Tadel *et al*., 2019).

The inverse solution was calculated using the weighted minimum norm estimation method and the measure standardized low resolution brain electromagnetic tomography as implemented in the Brainstorm software (Hamalainen & Ilmoniemi, 1994; Baillet *et al*., 2001; Pascual-Marqui, 2002). We chose to use a distributed source imaging model rather than a single dipole model since we expected multiple cortical regions to be modulated during feedback processing. Source activity (in the time domain) was estimated for sources with unconstrained orientation, that is source activity was calculated for three dipole orientations at each cortical location. The estimation of source activity via minimum norm estimators takes into account the level of noise in the sensors and hence requires an estimation of the noise in the recordings (Hauk, 2004). Thus, noise statistics (noise covariance across all sensors) were calculated across all trials from a pre-stimulus time, i.e. prior to information of the outcome (−2000 ms to 0 ms). Finally, we obtained source space time series at each cortical position.

### Time-frequency analyses in source space

Since cortical sources cannot be directly inferred from scalp time-frequency data, we computed time-frequency decomposition on the source time series using Morlet-wavelet analyses. Wavelets had a full-width at half-maximum of 3000 ms and a frequency of 1 Hz (corresponding to 7 cycles). Again, the analyses were constrained to the averaged theta band (4 – 8 Hz) and the averaged higher beta band (25 – 35 Hz).

In the first step of the analyses, these analyses were run on the full cortical map (entire cortical sheet) for all trials independent of the outcome to test our assumption that regions of the prefrontal cortex respond to outcome information in general. In support of this view, we observed strong oscillatory power over the SFG and RMFG suggesting that both regions engage in processing motor outcomes. Thus, in the following steps we constrained the analyses to SFG and RMFG. These were defined from pre-specified cortical areas of the Desikan-Killiany parcellation scheme which subdivides the cortex into gyral based ROI (Desikan *et al*., 2006).

Subsequently, time-frequency data of the ROI were computed for *S+^(n)^* and *S-^(n)^* trials. In our second analyses, we again faced the problem of differences in trial numbers between the different outcomes for every participant. This could result in meaningful differences in signal-to-noise ratios and bias the comparisons between the conditions (see also behavioural data analysis). Thus, we performed bootstrapping by sampling multiple new datasets with replacement. This resulted in 100×1,000 random trials per participant and condition. As for the behavioural analysis, trials were grouped according to the outcomes of the previous two trials in *S+^(n)^ & S+^(n-1)^, S+^(n)^ & S-^(n-1)^, S-^(n)^ & S+^(n-1)^* and *S-^(n)^ & S-^(n-1)^*. The averaged and normalised data were used for statistical analyses.

### Statistics on time-frequency data in source space

Statistical analyses were performed on time-frequency source time series in the pre-selected time window of interest (250 ms – 550 ms after outcome information). To reduce the number of tests, statistical analyses was performed on specified data points (in steps of approximately 50 ms, i.e. at 250 ms, 300 ms, … 550 ms, according to the sampling frequency). Further, we constrained our analyses to selected comparisons of interest. In the first step, we compared theta and high beta power time courses for all ROI between S+*^(n)^* and S-*^(n)^* trials. In the second step, we compared theta and high beta time courses for all ROI between outcomes i) *S+^(n)^ & S+^(n-1)^ and S+^(n)^ & S-^(n-1)^* and ii) *S-^(n)^ & S+^(n-1)^* and *S-^(n)^ & S-^(n-1)^*. In all cases, we tested whether the data fulfil the criteria for statistics on normally distributed data. For this purpose, we plotted histograms and performed Kolmogorov-Smirnov tests. Given that the data were not normally distributed, we performed Wilcoxon signed rank tests between all comparisons of interest. The FDR method was used to correct for multiple comparisons. Critical P-values are presented in the Results section, i.e. the adjusted significance threshold after correcting for the FDR. MATLAB R2019a (The MathWorks, Natick, Massachusetts, United States, R2019a) was used to compute all statistical analyses of the EEG data.

## Acknowledgements

P.W. was supported by the German Academic Exchange Service and University of Copenhagen. M.E.S. was supported by the Danish Ministry of Culture and Manufacturer Vilhelm Pedersen and his wife’s Memorial Scholarship. M.M.B. was supported the Danish Ministry of Culture. J.L.J. was supported by Nordea-fonden.

## Author contributions

P.W., M.E.S., C.R., M.M.B. and J.L.J. conceived and designed the research. P.W. performed the experiments. P.W. and C.R. analysed the data. P.W., M.E.S., C.R., M.M.B. and J.L.J interpreted the results of experiments. P.W. drafted the paper. P.W., M.E.S., C.R., M.M.B. and J.L.J. edited and revised the manuscript. All authors approved the final version of manuscript.

## Data and code availability

Data (Behaviour) and code (Behaviour and EEG) are available at https://data.mendeley.com (DOI: 10.17632/cw73pv9ct4.1). EEG data were not uploaded due to their large size but are available from the corresponding author on request.

