## Supplementary Information for "Behavioural variability and cortical electrophysiological signals depend on recent outcomes during human reinforcement motor learning"

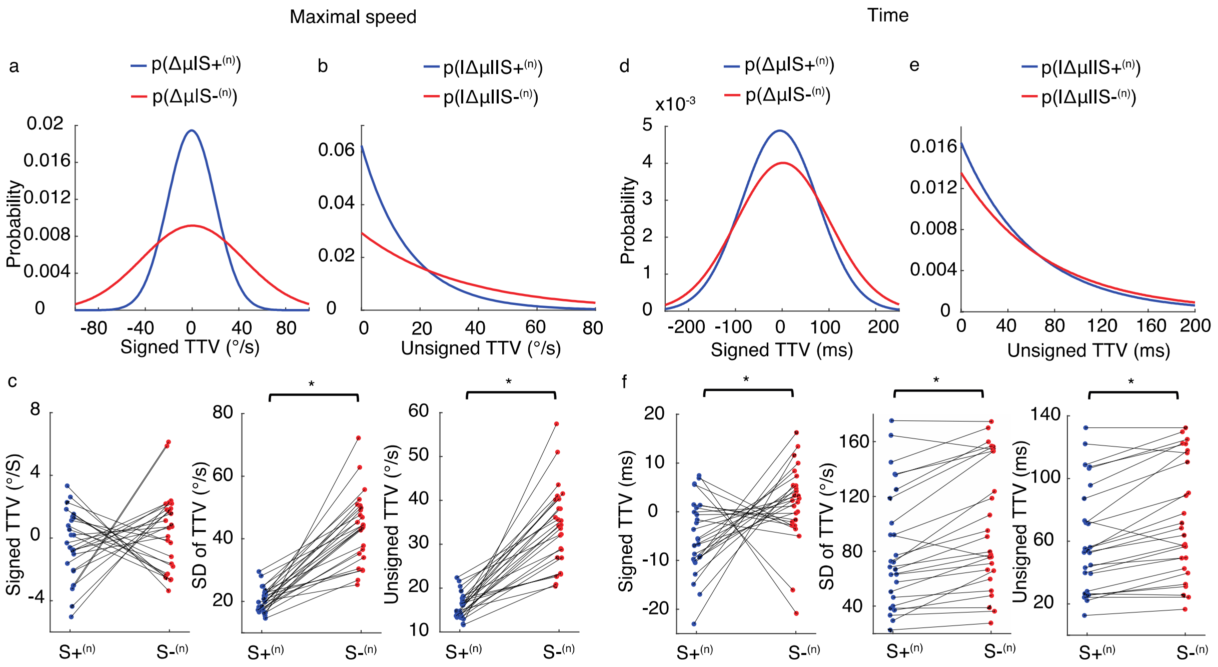


**Supplementary Figure 1. Reinforcement processes generalize to outcome-irrelevant parameters**

**a** The grand average normal distribution for signed changes in TTV of maximal movement speed after S+$^{(n)}$ and S-$^{(n)}$ trials. **b** The grand average exponential distribution for unsigned changes in TTV of maximal movement speed after S+$^{(n)}$ and S-$^{(n)}$ trials. **c** Individual differences in signed TTV (left panel), SD of TTV (middle panel) and unsigned TTV (right panel) of maximal movement speed after S+$^{(n)}$ and S-$^{(n)}$ trials**.** The analyses showed no significant differences between S+$^{(n)}$ and S-$^{(n)}$ trials in signed TTV (paired t-test, $t = -0.9$, $P = 0.335$) but in SD of TTV (F-test, $F = 108.1$, $P < 0.001$) and unsigned TTV (F-test, $F = 96.9$, $P < 0.001$). Note that x-axis values were jittered to more clearly present the data. **d** The grand average normal distribution for signed changes in TTV of movement time after S+$^{(n)}$ and S-$^{(n)}$ trials. **e** The grand average exponential distribution for unsigned changes in TTV of movement time after S+$^{(n)}$ and S-$^{(n)}$ trials. **f** Individual differences in signed TTV (left panel), SD of TTV (middle panel) and unsigned TTV (right panel) of movement time after S+$^{(n)}$ and S-$^{(n)}$ trials**.** The analyses showed no significant differences (after adjusting for multiple comparisons) between S+$^{(n)}$ and S-$^{(n)}$ trials in signed TTV (paired t-test, $t = -2.3$, $P = 0.029$) but in SD of TTV (F-test, $F = 28.2$, $P < 0.001$) and unsigned TTV (F-test, $F = 26.7$, $P < 0.001$). Note that x-axis values were jittered to more clearly present the data.

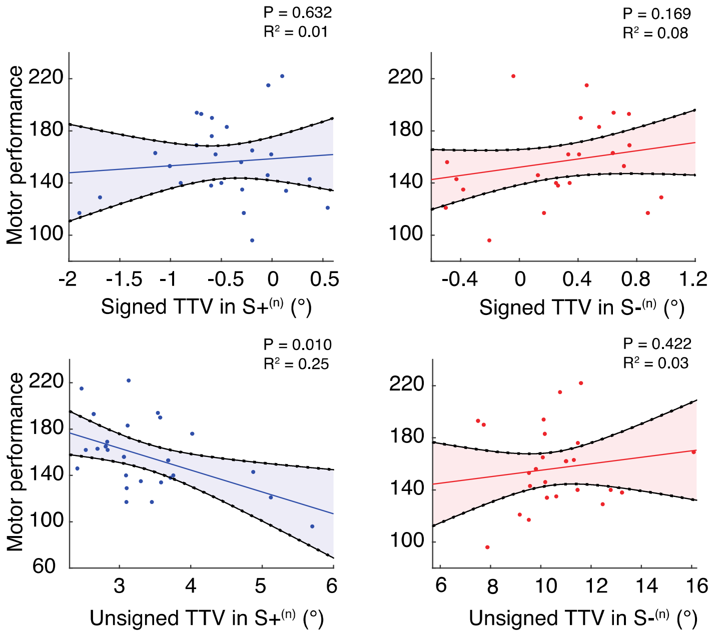


**Supplementary Figure 2. Changes in TTV after S+ outcomes predict overall motor performance**

Scatter plot showing the association between motor performance and signed TTV after S+$^{(n)}$ trials (linear regression: F = 0.2, P = 0.632, $R^2$ = 0.01, left top panel) and signed TTV after S-$^{(n)}$ trials (linear regression: F = 2.0, P = 0.169, $R^2$ = 0.08, right top panel), unsigned TTV after S+$^{(n)}$ trials (linear regression: F = 7.8, P = 0.010, $R^2$ = 0.25, left bottom panel) and unsigned TTV after S-$^{(n)}$ trials (linear regression: F = 0.7, P = 0.422, $R^2$ = 0.03, right bottom panel). Each dot represents an individual participant. The lines represent the fitted regression line. Shading is 95% confidence intervals of the regression.

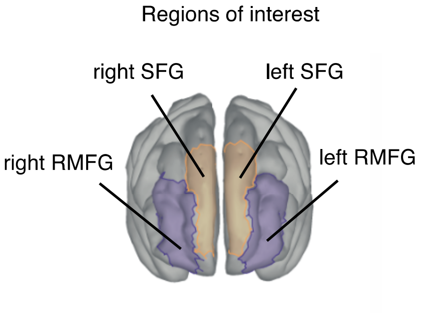


**Supplementary Figure 3. Cortical representation of the ROI**

These were the bilateral superior frontal gyrus (SFG) and bilateral rostral middle frontal gyrus (RMFG). The ROI were based on the Desikan-Killiany parcellation scheme which subdivides the cortex into gyral based ROI (Desikan *et al.*, 2006).

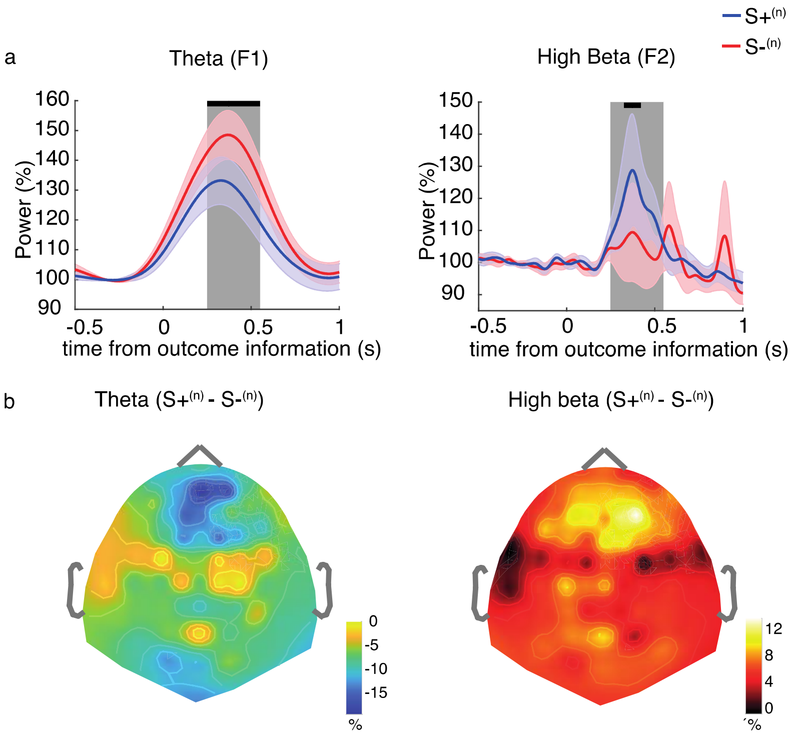


**Supplementary Figure 4. Oscillatory signals in frontal sensors depend on the previous outcome**
**a** Grand average power time courses for theta (left panel) and high beta (right panel) frequencies during outcome processing (-0.5 s – 1 s relative to outcome information) for S+$^{(n)}$ trials and S-$^{(n)}$ trials. Data are from channel F1 (theta) and F2 (high beta). The shaded rectangle highlights the time window of interest 250 ms – 550 ms. Shading around the mean represent the standard error of the mean. The black horizontal line indicates significant differences between the conditions (statistical tests were performed in steps of 50 ms). The critical alpha values were P = 0.037 for theta and P = 0.008 for high beta frequencies. **b** Topographical distribution of theta (left panel) and high beta (right panel) power during outcome processing (250 – 550 ms after outcome information). The data show the difference between S+$^{(n)}$ trials and S-$^{(n)}$ trials. Note that we used different colours to plot the topographical power distribution for theta and high beta frequencies since the data have different scales.

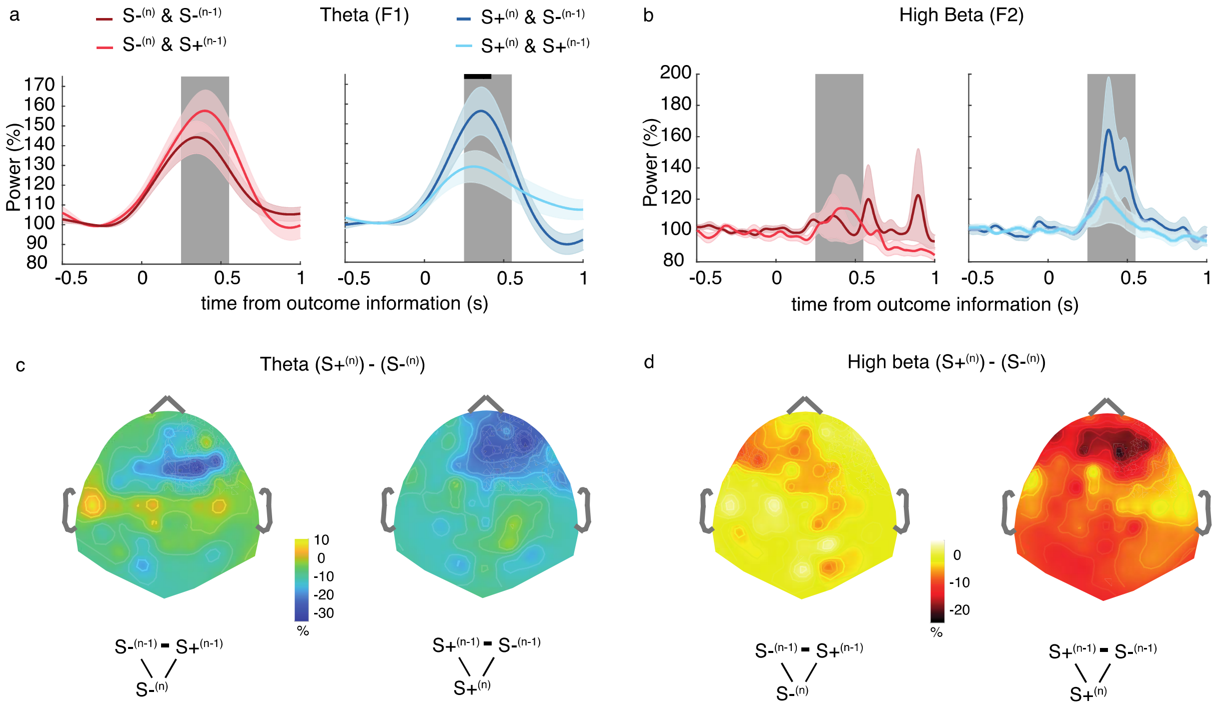


**Supplementary Figure 5. Oscillatory signals in frontal sensors depend on the outcomes of the past two movements**

**a & b** Grand average power time courses for theta (a) and high beta (b) frequencies during outcome processing (-0.5 s – 1 s relative to outcome information) for S-$^{(n)}$ & S-$^{(n-1)}$ trials, S-$^{(n)}$ & S+$^{(n-1)}$ trials, S-$^{+n)}$ & S-$^{(n-1)}$ trials and S+$^{(n)}$ & S+$^{(n-1)}$ trials. Data are from channel F1 (theta) and F2 (high beta). The shaded rectangle highlights the time window of interest 250 ms – 550 ms. Shading around the mean represent the standard error of the mean. The black horizontal line indicates significant differences between the conditions (statistical tests were performed in steps of 50 ms). The critical alpha values were P = 0.023 for theta frequencies (contrast: S-$^{+n)}$ & S-$^{(n-1)}$ trials and S+$^{(n)}$ & S+$^{(n-1)}$ trials). All other comparisons did not reach statistical significance. **c & d** Topographical distribution of theta (c) and high beta (d) power during outcome processing (250 – 550 ms after outcome information). The data show the difference between the conditions of interest. Note that we used different colours to plot the topographical power distribution for theta and high beta frequencies since the data have different scales.
